# Human Choroid Plexus Organoids Progress to Postnatal-like Maturation and Reveal a Closing Window of Developmental Plasticity

**DOI:** 10.64898/2026.10.12.758506

**Authors:** Elizabeth J. Apsley, See Swee Tang, Elisa Vitiello, Ivan Imaz-Rosshandler, Marian Fernandez-Otero, Kyriaki Sapidou, Sean A. Burnap, Weston B. Struwe, Milos Pjanic, John F. Fullard, Jaroslav Bendl, Panos Roussos, Katherine R. Long, Fanni Gergely, Madeline A. Lancaster, Laura Pellegrini

**Affiliations:** Centre for Developmental Neurobiology, Institute of Psychiatry, Psychology & Neuroscience, King’s College London, London SE1 1UL, UK; Department of Biochemistry, University of Oxford, Parks Rd, Oxford OX1 3QU, UK; Altos Lab, San Diego, CA, 92121, USA; Kavli Institute for Nanoscience Discovery, Dorothy Crowfoot Hodgkin Building, University of Oxford, OX1 3QU, United Kingdom; Center for Disease Neurogenomics, Icahn School of Medicine at Mount Sinai, New York, NY 10029, USA; Friedman Brain Institute, Icahn School of Medicine at Mount Sinai, New York, NY 10029, USA; Department of Psychiatry, Icahn School of Medicine at Mount Sinai, New York, NY 10029, USA; Department of Genetics and Genomic Science, Icahn School of Medicine at Mount Sinai, New York, NY, 10029, USA; Mental Illness Research, Education and Clinical Center VISN2, James J. Peters VA Medical Center, Bronx, NY, 10468, USA; Center for Precision Medicine and Translational Therapeutics, James J. Peters VA Medical Center, Bronx, NY, 10468, USA; MRC Centre for Neurodevelopmental Disorders, King’s College London, London, UK; MRC Laboratory of Molecular Biology, Cambridge Biomedical Campus, Francis Crick Avenue, Cambridge CB2 0QH, UK

## Abstract

The choroid plexus (ChP) produces cerebrospinal fluid (CSF) and forms the blood-CSF barrier, yet the developmental programme driving its maturation in humans remains poorly understood. Using a 15-timepoint single-cell RNA sequencing atlas of human ChP organoids benchmarked against human tissue, we resolve an ordered differentiation process comprising cell cycle exit, ciliogenesis and ion transporter expression. Maturation culminates in a transcriptional state with features of human adult ChP, establishing the first *in vitro* ChP model to reach postnatal-like milestones. Centrosome proteomics identified previously uncharacterised, stage-specific regulators of ciliogenesis. To probe developmental plasticity, we perturbed ChP organoids with valproic acid (VPA), a clinically used antiepileptic and established teratogen. Early VPA-mediated disruption of multiciliogenesis and secretion recovered upon drug withdrawal, whereas late treatment washout resulted in persistent impairment of mature transporters. Together, these findings resolve the developmental programme by which the ChP reaches functionally mature identity and identify a stage-dependent plasticity window during its maturation.

## Introduction

The choroid plexus (ChP) is the main source of cerebrospinal fluid (CSF) and forms an essential brain barrier, separating the CSF from the systemic circulation. Located in each ventricle of the brain, this sheet-like structure is composed of a polarised multi-ciliated epithelium surrounding a stromal compartment with fibroblasts, immune cells and fenestrated capillaries ^1–4^. Beyond its secretory and barrier functions, the ChP is increasingly recognised as a central regulator of brain homeostasis, coordinating neural progenitor activity during development ^5,6^ and a key immunological hub, modulating neuroinflammatory responses throughout life ^7,8^.

The ChP is among the earliest structures to form during brain development. In humans, the lateral ventricle ChP emerges from the cortical hem approximately by 7 gestation weeks and undergoes terminal differentiation before birth, a remarkably rapid developmental trajectory compared to other regions in the brain, such as the cerebral cortex ^9–11^. Unlike rapidly renewing epithelia such as the gut or respiratory tract, the adult ChP lacks a dedicated stem cell niche, and cross-species studies suggest that mature ChP epithelial cells display limited proliferation ^12–15^. *In vitro* models of adult mouse ChP epithelial cells show capability of re-entering the cell cycle and proliferating in response to specific growth factors like EGF, IGF-1, and IGF-II signalling ^12^. Another central feature of ChP differentiation is the acquisition of multiple apical nodal (9+0) cilia, critical for mechanosensation and signalling, which regress postnatally ^1,16,17^.

Despite the central role of the ChP in brain development, human ChP biology remains largely inferred from mouse studies and histological observations ^10,11^. As such, fundamental questions about how the human ChP epithelium acquires its mature identity remain unanswered. Human ChP organoids provide a tractable system to address these questions: they recapitulate the polarised epithelial monolayer cytoarchitecture, ciliation, selective barrier and secretory function with CSF-like fluid production ^18^. However, whether organoids can progress beyond their foetal-like state to acquire adult transcriptional and functional ChP identity has not been established.

In this study, we generate a 15-time point single-cell RNA sequencing (scRNA-seq) atlas of human ChP organoids from day 14 to day 130 of differentiation, benchmarked against foetal and adult human ChP tissue. We show that ChP epithelial maturation proceeds through sequential, ordered phases (proliferation exit, multiciliogenesis and secretory maturation). ChP epithelial cells acquire transcriptional and functional profiles similar to adult human ChP, establishing for the first time an *in vitro* model that shares features of adult ChP. Using centrosome affinity capture mass spectrometry (CAPtureMS), we identify stage-specific cilia regulators and novel centrosome associated proteins with no prior, or poor, characterisation in ciliogenesis, but with a strong temporal pattern matching the cilia remodelling across ChP. Finally, to test whether the ordered maturation programme retains the capacity to recover from perturbation applied during defined developmental windows, we use valproic acid (VPA), a clinically used HDAC inhibitor previously used as a pharmacological challenge for ChP ^19^. We show that VPA-mediated disruption of multiciliogenesis is reversible upon withdrawal. We also observed disruption of CSF secretion and downregulation of ion transporter SLC5A5 with VPA which were reversible upon withdrawal only at early developmental stages but persisted during the late secretory phase, defining a window of developmental plasticity. Together, these findings identify the maturation programme of human ChP epithelium as well as the developmental window during which the ChP remains plastic, with implications for understanding the timing-dependent consequences of teratogen exposure during human brain development.

## Results

### A longitudinal single-cell atlas resolves the ordered differentiation program of human ChP organoid development

To characterise the cellular composition and developmental trajectory of the human ChP, we used our previously established ChP organoid model ^18^. These organoids secrete CSF-like fluid (iCSF) and form a functional barrier with selective permeability. ChP organoids also recapitulate key cytoarchitecture of the human foetal ChP including a polarised TTR-positive epithelial monolayer with multiciliated cells (Fig.1A).

**Figure 1.**
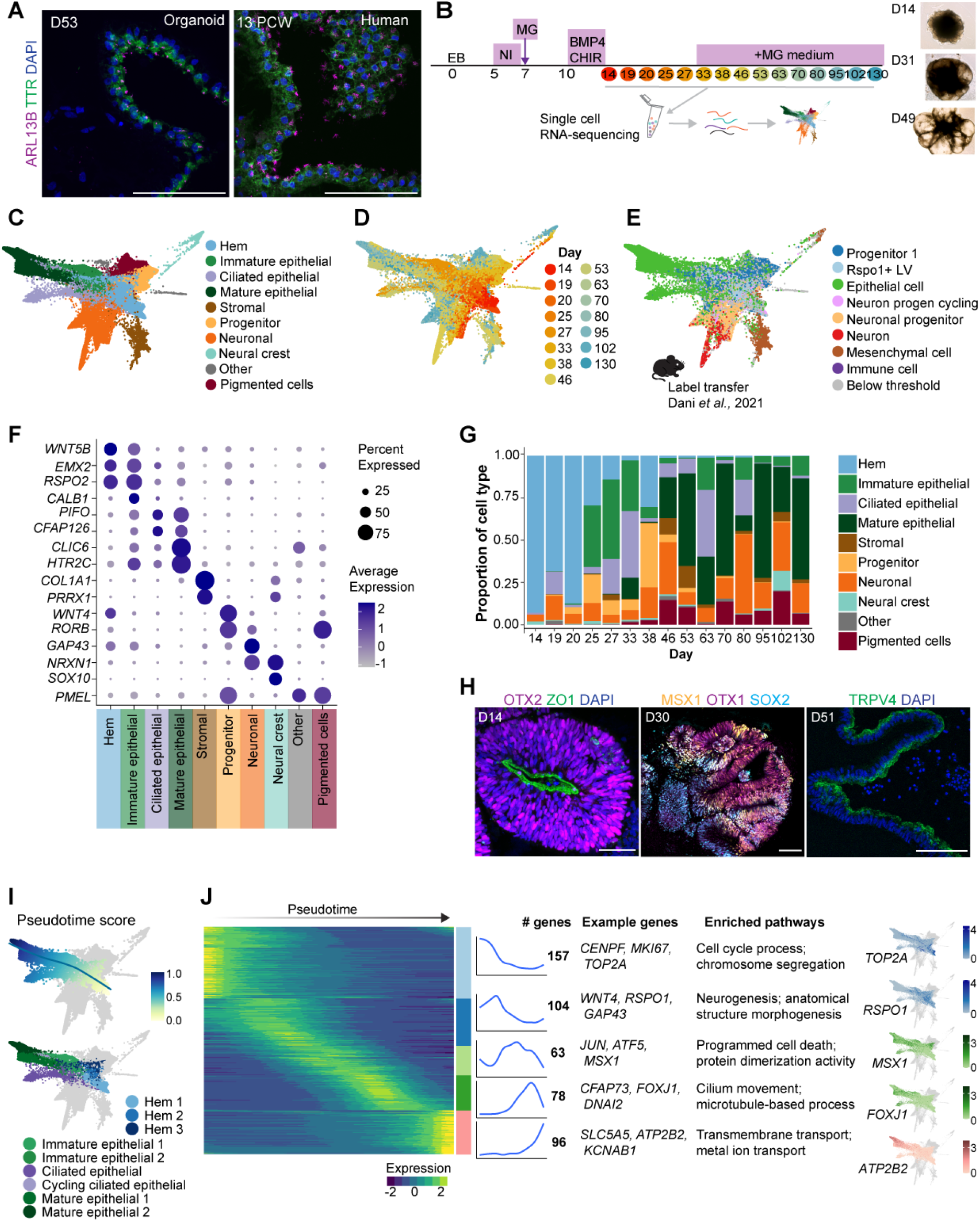
A longitudinal single-cell atlas of human ChP organoid development identifies diverse cell lineages reflecting in vivo composition. **(A)** Ciliated ChP epithelium in D53 organoid and 13pcw human fetal tissue immunostained for TTR (green) and ARL13B (cilia, magenta). Scale bar: 100 µm. **(B)** Schematic of ChP organoid generation and sample collection for scRNA-seq. Representative images of organoids showing neuroepithelial buds at D14, early epithelial cyst formation (D31) and later ChP cyst (D49). EB, embryoid body; NI, neural induction; MG, matrigel. **(C)** FA plot labelled by broad cell type cluster annotation showing the diversity of cell types present in ChP organoids. **(D)** FA plot labelled by day of sample collection (D14 to D130) displaying how cell populations change over differentiation. **(E)** FA plot coloured by label transfer of cell type annotations from embryonic mouse lateral ventricle ChP scRNA-seq ^20^. **(F)** Expression of selected markers distinguishing broad ChP cell types. Dot size indicates the percentage of cells in each cluster expressing the gene and colour scale shows average gene expression. **(G)** Proportion of cell types present at each sample age in ChP organoid scRNA-sequencing. **(H)** Organoid sections across time show differentiation from cortical hem to ChP epithelium. Neuroepithelial buds present in D14 ChP organoids formed from early cortical hem (OTX2, magenta), with apical tight junctions (ZO1, green). Scale bar: 50µm. D30 organoids contain cortical hem and neuronal progenitors labelled by MSX1 (yellow), OTX1 (magenta) and SOX2 (cyan). Scale bar: 100µm. D51 organoids formed polarised epithelium (TRPV4, green). Scale bar: 100µm. **(I)** Pseudotime score and trajectory for cells in epithelial lineage shown on FA plot. Epithelial lineage cell type clusters included in epithelial lineage trajectory shown on FA plot. **(J)** Differentially expressed genes across epithelial lineage development. Heatmap shows gene expression over pseudotime, grouped into modules by shared expression patterns as indicated by sidebar. Further characteristics of each module are shown on right: line graphs of average expression trend over pseudotime, number (#) of genes, top enriched GO terms, and example genes for each module (Table S3, S4). Nuclei are stained with DAPI (blue) throughout.

To resolve cellular dynamics across human ChP maturation, we generated a longitudinal single-cell transcriptomic atlas of ChP organoids from day 14 to day 130 of differentiation (47,697 cells post-QC), integrating a previously published dataset (Fig.1B-D, Fig.S1A-G, Table S1) ^18^. Unsupervised clustering identified 21 cell populations, which were annotated based on specific marker expression and grouped to 10 broad cell types (Fig. 1C, 1F, Fig. S1D-G, Table S2). Organoid cell types could be classified into three major compartments: epithelial, stromal, and neural. The epithelial compartments included cortical hem, a Wnt signalling centre involved in ChP patterning (*WNT4*, *WNT5B, EMX2, RSPO2*), ChP epithelial cells (*CLIC6, HTR2C*), and ChP ciliated epithelial cells (*PIFO, CFAP126)*. The stromal compartment was defined by fibroblast markers (*COL1A1, PRRX1, DCN, LUM*). The neural compartment comprised neuronal progenitors (*EPHB1*, *PAX6*) and neurons (*RORB*, *GAP43, NRXN1*) (Fig.1C, 1F, Table S2). We additionally identified neural crest (*SOX10*) and two clusters expressing genes related to pigment production consistent with a neural crest-derived pigmented cells (*PMEL*, *DCT*) (Fig.1C, 1F, Fig.S1G). We validated cluster annotations by label transfer analysis using a published scRNAseq data from embryonic mouse ChP ^20^ and correlation of gene expression with a recent 2D human ChP model ^17^, both of which confirmed cell type assignments (Fig. 1E, Fig. S1H). We previously described that ChP organoids recapitulate two morphologically and ultrastructurally distinct ChP epithelial subpopulations: cilia-rich “light” cells and mitochondria-rich “dark” cells ^18,21^. These two states have been also characterised in a recent 2D *in vitro* model of ChP as metabolic (type 1) and ciliogenic (type 2) epithelial cells ^17^. In our atlas, early ciliated epithelial cells aligned with the 2D ciliogenic/type 2 population or “light” cells (*FOXJ1, CCNO*), while more mature epithelial clusters (Mature epithelial 1 and 2), enriched for transmembrane transporters (*ATP1B2* and *FOLR1),* corresponded to the 2D metabolic/type 1 “dark” population (Fig. S1H-I), indicating that organoids capture both ciliogenic and metabolic axes of ChP epithelial identity.

Temporal analysis of cluster proportions revealed a shift from cortical hem to epithelial identity between day 28 and day 35, coinciding with morphological emergence of CSF-filled cysts (Fig. 1G). Early organoids express ChP progenitor transcription factors (OTX1, OTX2, MSX1), while differentiated ChP epithelial cells express canonical markers such as transient receptor potential, vanilloid 4 (TRPV4) (Fig. 1H). This temporal sequence mirrors the *in vivo* human lateral ventricle ChP epithelial specification from cortical hem, which occurs by approximately 6-7 post-conceptional weeks (pcw) ^3,9,10^.

To identify the transcriptional programmes underlying this developmental trajectory, we performed pseudotime analysis focused on the cortical hem and epithelial lineage (Fig. 1I). Following the trajectory from early progenitors to mature epithelial cells, pseudotime correlated with sample chronological age up to day 70 before plateauing at later timepoints (Fig. S1J). Differential expression analysis along pseudotime identified five gene modules (Fig. 1J, Table S3). Two early modules of genes associated with cell division (*CENPF, MKI67, TOP2A*) and neurogenesis (*WNT4, RSPO1, GAP43*), prevalent in cortical hem clusters, decreased sharply during epithelial differentiation (after day 33), indicating exit from cell cycle and loss of neural lineage specification (Fig. 1J, Fig. S2C, Table S4). Two late modules captured epithelial maturation: a ciliogenesis module (*CFAP73, FOXJ1, DNAI2*), peaking between days 40 and 60 before declining, and a transmembrane transport/secretory module (*SLC5A5, ATP2B2, KCNAB1*), progressively increasing (Fig. 1J, Table S4). Among transcription factors, ChP epithelial differentiation was driven mainly by loss of neuroepithelial and early cortical regulators (*FOXG1, EMX2, PAX3, HES1*), with later modules containing the ciliogenesis regulator FOXJ1 (module 4) and the immune and vascular regulators *POU2AF1* and *EPAS1* (module 5) (Fig. S1L). Together, these modules describe an ordered differentiation programme in which proliferative and neurogenic potential are declining in parallel, followed sequentially by ciliogenesis and secretory maturation. We next examined each of these phases, beginning with the kinetics of proliferative exit.

### ChP epithelial cells reach terminal differentiation *in vitro*

We first validated the earliest feature of the epithelial lineage trajectory: the rapid downregulation of cell-cycle genes. Cell cycle scoring showed a progressive reduction of cells in G2M phase, reaching 2% of all organoid cells at late timepoints (Fig. 2A, Fig. S2A), in agreement with limited observations from human foetal ChP *in vivo* ^14,17^. To confirm cell-cycle exit, we used a FUCCI (Fluorescent Ubiquitination-based Cell Cycle Indicator) reporter line ^22^ (Fig. 2B). ChP epithelial cells shifted to predominantly G1/G0 by day 35, with few S/G2/M cells visible by day 70 (Fig. 2B). Flow cytometry analysis quantified the same switch, with only 3.1% S/G2/M cells present at day 80 (Fig. 2C). Together, these results demonstrate that ChP organoids recapitulate the rapid proliferative decline of human ChP epithelium, transitioning to a predominantly post-mitotic monolayer by day 70.

**Figure 2.**
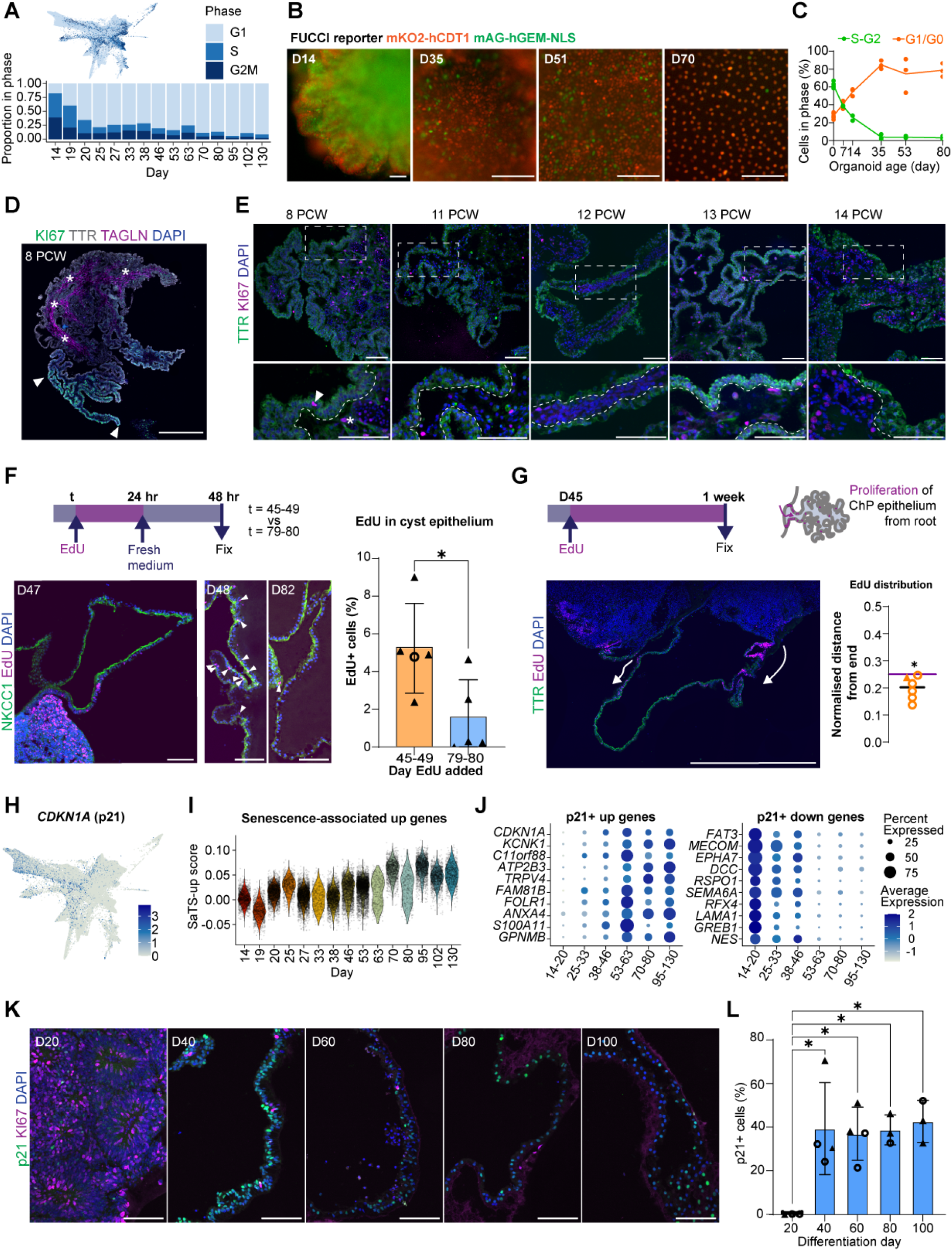
ChP epithelial cells reach terminal differentiation *in vitro* and recapitulate *in vivo* conveyor-belt dynamics. **(A)** Predicted cell cycle phase in ChP organoid scRNA-seq shown on FA plot and proportion per phase across time. **(B)** Live cell images of H9-FUCCI cell-cycle reporter line D14-D70. Reporter labels G1/G0 in orange (mKO2-CDT1) and S/G2 in green (mAG-GEM). Images from D35 onwards focus on organoid cysts. Scale bar: 100µm. **(C)** Percentage of cells in G1/G0 and in S/G2 quantified by flow cytometry across ChP organoid differentiation. Multiple organoids were pooled for each sample (n=3). **(D)** Human fetal ChP at 8pcw showing heterogeneous cell division. In localised region (arrows) epithelial cells (TTR, grey) are highly proliferative (KI67, green), whilst proliferation elsewhere is largely limited to stromal regions (star, TAGLN, magenta). Scale bar: 500µm. **(E)** Progressive decline of proliferation (KI67, magenta) in ChP epithelium (TTR, green) across development, in human fetal ChP sections from 8 to 14 pcw. Dashed lines separate epithelial from stromal layers. Example of proliferative cells in epithelium (arrow) and stroma (star). Scale bar: 100µm. **(F)** Confocal images of organoids labelled for epithelial cells (NKCC1, green) and EdU (magenta) with EdU added at mid (D45-49) and later (D79-80) stages of differentiation. Percentage of EdU+ cells was quantified in NKCC1+ cyst epithelium (n=5). Scale bar: 100µm. Significant difference in average EdU frequency between early and later time points, t-test * p < 0.05. **(G)** Distribution of proliferation across organoid cyst after 1 week of 1uM EdU exposure. Relative distance of EdU+ cells measured from the nearest end. Points represent average distances per organoid (n= 6). Representative image, labelled for TTR (green), EdU (magenta) and nuclei (dapi, blue). Scale bar: 1mm. **(H)** Expression of cyclin-dependent kinase inhibitor *CDKN1A* (p21) in ChP organoid scRNA-seq. **(I)** Module score for senescence associated transcriptomic signature (SaTS) upregulated genes ^24^ across differentiation time in epithelial lineage cells. **(J)** Top differentially expressed genes between p21+ and p21-cells within the ChP epithelial lineage. Dot size indicates the percentage of cells in each time period expressing the gene, colour scale shows average gene expression. **(K)** ChP organoids collected across time, immunostained for p21 (green) and KI67 (magenta). Scale bar: 100 µm. **(L)** Quantification of p21-positive epithelial cells over time in ChP differentiation. Points represent average values for organoids (n = 3-4). ANOVA followed by Tukey HSD, * p<0.05. Nuclei are stained with DAPI (blue) throughout. “N” indicates independent organoid batches. Cell lines are indicated by point shape in plots: solid triangle H1, open circle H9.

To benchmark these dynamics against the developing ChP *in vivo*, we examined Ki67 expression in human foetal tissue across a developmental time course (Fig. 2D). At 8pcw, epithelial proliferation was spatially heterogeneous with Ki67-positive cells concentrated near the root of the plexus (Fig. 2D). We observed a decline in epithelial proliferation by 11 to 14pcw, with Ki67-positive cells mostly present in the stroma (Fig. 2E). Although this does not exclude proliferation still being present at the root of the ChP at later time points.

To resolve the spatial organisation of proliferation along the epithelial monolayer, we performed EdU incorporation assays. EdU+ cells were predominantly located in the core of ChP organoids (Fig. 2F). Among NKCC1-positive epithelial cells, the proportion of EdU+ cells decreased from 5.2% at day 45-49 to 1.5% at day 79-80 (Fig. 2F). Tracking EdU+ cells over a week revealed that epithelial proliferative cells were predominantly located towards the root of the organoid. This is consistent with the conveyor-belt model of ChP expansion described *in vivo* and with a low-frequency, spatially distinct proliferative pool actively maintained to drive tissue expansion (Fig. 2G) ^1,15,23^.

Accompanying cell cycle exit, recent work has identified a non-inflammatory senescent state emerging in ChP epithelial cells during mouse embryonic development and persisting into adulthood ^24^. To examine whether this senescent state occurs in human ChP development we analysed expression of *CDKN1A* (p21), hallmark of cellular senescence. *CDKN1A* expression increased over time in the epithelial lineage (Fig. 2H), whilst the alternative regulator of senescence *CDKN2A* (p16) remained low, consistent with mouse findings ^24^ (Fig. S2B). Development of a senescent state may explain GO-terms related to cellular stress enriched in epithelial lineage module three. This gene module contained *CDKN1C* (p57) another cyclin-dependent kinase inhibitor associated with senescence ^25^ (Fig. S2B). We next calculated module scores for previously defined senescence-associated transcriptomic signatures (SaTS) ^24^. Across time the organoid epithelial lineage showed increased expression of SaTS upregulated genes and decreased expression of SaTS downregulated genes, providing further evidence for development of a senescent state (Fig. 2I, Fig. S2C). Comparison of p21+ to p21-cells within the epithelial lineage confirmed downregulation of genes associated with cell cycle, neuronal progenitors and anabolic metabolism (Fig. 2J, Fig. S2D). Immunostaining verified presence of p21+ cells in ChP epithelium by day 40 and confirmed absence of Ki67-staining in these cells (Fig. 2K, L). Genes upregulated in p21+ cells related to respiration, cilia, and the core ion transport, calcium signalling and folate receptor machinery required for active CSF production and barrier function (Fig. 2J, Fig. S2D). These findings are in accordance with the proposed long-term non-degenerative senescence state supporting ChP function and further demonstrate that organoids recapitulate terminal differentiation and functional specialisation of mature ChP epithelial cells.

### Temporal remodelling of ChP epithelial cilia in organoids

The pseudotime analysis of the epithelial lineage identified a transient peak in genes related to ciliogenesis (Fig. 1J). Enrichment was confirmed using an independent list of genes associated with multi-ciliated cells ^26^ (Fig. 3A). Key genes, including master ciliogenesis regulator *GMNC*, ciliogenesis-inducing transcription factor *FOXJ1*, and cilia components *DNAI2* and *CFAP73*, peaked between days 30 and 60 (Fig 3B). Immunostaining for TP73, transcription regulator of multiciliogenesis ^27^, further demonstrates the peak of ciliogenesis around day 40 of organoid development (Fig. 3C).

**Figure 3.**
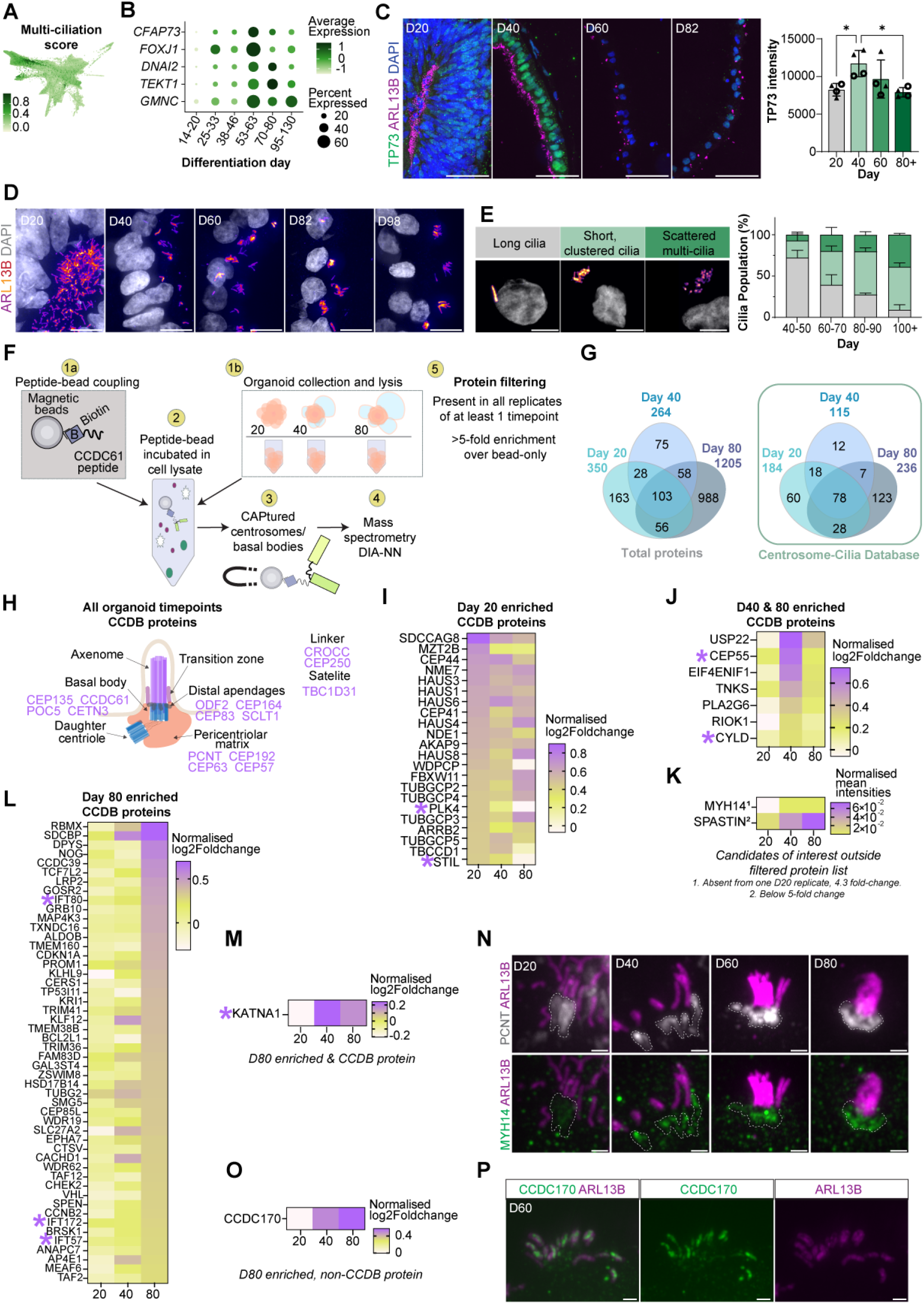
Centrosome proteomics identifies novel regulators of ChP ciliogenesis. **(A)** Multi-ciliation gene score from Lewis and Stracker ^26^ projected on FA plot. **(B)** Dotplot of cilia gene expression in epithelial lineage cells over time. Dot size indicates the percentage of cells expressing the gene, colour scale shows average gene expression scaled per gene. **(C)** TP73 expression peaks in developing ChP epithelium. Organoids across differentiation immunostained for TP73 (green) and ARL13B (magenta). Scale bar: 100µm. Quantification of TP73 intensity over time, points represent organoids from separate differentiations (n=4), Cell lines are indicated by point shape: solid triangle H1, open circle H9. ANOVA followed by Tukey HSD, * p <0.05. **(D)** ChP epithelial cilia (ARL13B, fire) morphology across differentiation in immunostained organoid sections. Scale bar: 10µm. **(E)** ChP organoid cilia classified according to morphology: long cilia (grey), short, clustered cilia (light green) and scattered multi-cilia (dark green). Representative ARL13B-labelled images for each class, nuclei labelled with dapi (grey). Proportion of cilia per class over time, plotted as mean with standard error bars. Two-way ANOVA and Tukey multiple-comparison test, (n=4-5). Long cilia significantly decreases over time (adj. p < 0.0001 for D40-50 vs all other ages and D80-90 vs D100+; adj. p =< 0.01), with corresponding increases in clustered cilia (adj. p < 0.001 D40-50 vs all other ages) and scattered cilia (adj. p < 0.05 D40-50 vs D60-70 and D80-90; adj. p < 0.0001 D40-50 vs D100+; adj. p < 0.001 D60-70 vs D100+; adj. p < 0.01 D80-90 vs D100+). **(F)** Schematic of CAPture-MS proteomics workflow. Organoid samples collected at D20, D40 and D80. **(G)** Venn diagrams of proteins passing filtering criteria: full protein list (left) and within the centrosome-cilia database (CCDB, right) (Table S6). **(H)** Diagram of ciliary architecture and representative proteins identified at all three timepoints. **(I)** Heatmap of D20-only enriched proteins (log2foldchange compared to bead-only > 2.3, normalised to CEP135). Colour scale from low (yellow) to high (purple). **(J)** Heatmap of proteins enriched at both D40 and D80 (same filtering and colour scale as in I). **(K)** CAPture-MS intensity of MYH14 and Spastin across organoid timepoints. Values shown as log₂-fold change over bead-only, normalised to CEP135. **(L)** Heatmap of D80-only enriched CCDB annotated proteins D80 (same filtering and colour scale as in I). **(M)** CAPture-MS intensity of KATNA1 across organoid timepoints (same filtering and colour scale as in I). **(N)** Cilia complexes across D20 to D80 organoid development. Immunostaining for ARL13B (magenta) with PCNT (grey, top) or MYH14 (green, bottom). Dashed lines mark PCNT+ basal body regions. Scale bar: 1µm. **(O)** CAPture-MS intensity of CCDC170 across organoid timepoints (same filtering and colour scale as in I). **(P)** Representative images of CCDC170+ cilia complexes (green) at day D40 and D60 with cilia labelled by ARL13B (green) and nuclei by dapi (blue). Scale bar: 1µm.

Beyond the expression of ciliogenesis-associated genes, the morphology of ChP cilia changes across development. ChP epithelial cells transition from mono-ciliated to multi-ciliated during embryonic stages, before undergoing cilia regression postnatally resulting in more sparse ciliated cells ^1^. The spacing of cilia in multi-ciliated ChP cells has also been reported to change over time towards more scattered cilia in adult mice and humans ^16,17^. To assess these changes, we first quantified ARL13B-labelled cilium length in ChP organoids across multiple timepoints (Fig. 3D). Cilia length decreased, reaching a plateau of ∼1.7µm after day 60-70 (Fig. S3A), comparable with human postnatal cilia length (Ho et al., 2023). To better capture cilia heterogeneity, we applied a previously described classification distinguishing long, short and clustered, and scattered multi-cilia ^16^ (Fig. 3E). Quantification of these categories revealed a progressive shift from long, to shorter cilia in multi-cilia clumps or more scattered (Fig. 3E). In addition, we noted a difference in the range of cilia length across categories of cilia (long cilia: 2.52-2.99µm; short and clustered: 1.07-1.87µm; and scattered multi-cilia: 0.41-0.85µm), consistent with remodelling of ChP cilia towards mature conformation.

To investigate the centrosomal machinery underlying this remodelling, we profiled the centrosomal proteome across maturation. We collected organoids at day 20, 40, and 80 and performed centrosome affinity capture mass spectrometry (CAPture-MS) (Fig. 3F) ^28^. CAPture-MS analysis identified a total of 7080 proteins in d20, 8890 in d40 and 8493 in d80 organoids. PCA across timepoints revealed clear separation of samples by developmental stage (Fig. S3B). This continuous trajectory of proteomic change during ChP maturation further indicates robustness of the differentiation protocol. H1- and H9-derived day 40 organoids showed no significant differences in protein abundance and were pooled for subsequent analyses (Fig. S3C). GO enrichment analysis of CAPture-MS data across the three timepoints revealed consistent enrichment of centrosome- and cilia-related biological processes (Fig. S3D), confirming specificity of the pull-down. To verify the efficacy of CAPture protocol on organoids, filtered candidates were first intersected with the centrosomal protein list characterised in HEK-293 cells ^28^ (Table S5).

Identified proteins were then filtered in the following manner: candidates were included only if they were i) present in all CAPture replicates for specific stage, and ii) showed a five-fold enrichment in intensity relative to the corresponding bead-only samples (Fig. 3F). A fold-enrichment cutoff was necessary because D20 and D80 organoids had only one bead-only control, preventing standard statistical analysis. This identified 350, 264, and 1205 candidate proteins for D20, D40, and D80, respectively (Fig. 3G). Next, filtered candidates were intersected with the Centrosome-Cilia Database (CCDB), incorporating SysCilia Gold Standard and CentrosomeDB datasets ^29–32^ revealing significant overlap with known centrosome and cilia proteins (Table S6). Across all organoids, 326 CCDB proteins were identified, including stage-specific sets at D20 (184), D40 (115), and D80 (236) (Fig. 3G).

Among the proteins found in all timepoints, we detected several known centrosomal and basal body proteins, such as pericentriolar matrix proteins (PCNT, CEP192, CEP63), core centriolar structural proteins (CEP135, POC5, CETN3) and distal appendage proteins (CEP164, CEP83, SCLT1) (Fig 3H). Recovery of canonical centrosomal proteins across all timepoints confirmed the specificity of the CAPture pull-down. To identify candidates associated with ChP epithelial maturation, we compared filtered candidates across time points. The centriole duplication regulators PLK4 and STIL ^33^ were enriched at D20 and decreased progressively through D40 and D80 (Fig. 3I), consistent with loss of cycling cells during differentiation. Conversely, two established regulators of ciliary dynamics were enriched at D40 and D80 (Fig. 3J): CEP55, which recruits Aurora A kinase to centrosomes to promote ciliary disassembly ^34,35^, and CYLD, which regulates HDAC6-mediated axonemal tubulin acetylation and axoneme length ^36^ (Fig. 3J). Their coordinated late enrichment is consistent with the progressive cilia shortening observed during multiciliated cell maturation.

As a second validation, we examined Spastin, a microtubule-severing enzyme involved in cilia shortening and previously reported to increase during mouse ChP epithelial maturation ^1^. Spastin showed a progressive enrichment across differentiation (Fig. 3K), consistent with its reported dynamics *in vivo* and with the reduction in cilium length observed during organoid maturation. This trend extended to other cilia-related candidates enriched at D80, including the related microtubule-severing enzyme Katanin (KATNA1) ^37,38^, and multiple intraflagellar transport components required for cilium length maintenance, including IFT80, IFT172 and IFT57 (Fig. 3L, M, Table S6). Together these candidates reflect the coordinated acquisition of mature ciliary structural, transport and motility machinery during ChP maturation.

Among the D80-enriched candidates, MYH14, a non-muscle myosin linked to hereditary hearing loss and stereocilia organisation ^39–41^ and reported to localise near the basal bodies of multiciliated ependymal cells ^42^, emerged as a compelling candidate. MYH14 exhibited 4.43 log₂-fold enrichment over bead-only control at D20 and progressively increased in abundance across differentiation (Fig. 3K). Immunostaining confirmed MYH14 localisation at PCNT-labelled basal bodies, with increased relative recruitment during ChP maturation, consistent with a role in basal body remodelling and mature ciliary architecture (Fig. 3N).

Finally, we investigated D80-enriched candidates absent from the CCDB but with potential ciliary roles that, to our knowledge, have not been previously characterised in ChP. CCDC170 emerged as an intriguing candidate, as it localises to the base of primary cilia in IMCD3 cells and zebrafish embryos ^43^ and contributes to microtubule stabilisation ^44^. In our CAPture proteomics, CCDC170 showed progressive enrichment across differentiation (Fig. 3O) and co-immunostaining with the ARL13B confirmed specific CCDC170 localisation on ChP cilia in around 10% of cilia complexes (Fig. 3P). CCDC170 presence was variable across ciliated cells from D40 onwards, however the frequency trend peaked around D60 (Fig. S3K, L).

Together, these observations indicate that ChP cilia progressively remodel toward their mature postnatal morphology during organoid development and identify CCDC170 and MYH14 as candidate regulators that deserve further investigation.

### Organoids acquire mature transcriptional and functional ChP signatures

Acquisition of a mature secretory programme is the defining functional endpoint of ChP epithelial differentiation. To characterise this transition, we next examined the gene module associated transmembrane transport that increased progressively across maturation (Fig. 2C, Fig. 4A). Our epithelial trajectory analysis highlighted the temporal upregulation of genes associated with transmembrane transport of ions, including voltage-gated potassium channels (*KCNAB1, KCND3, KCNH5, KCNMA1*), voltage-gated calcium and sodium channels (*CACNA1C, CACNB2, SCN2A*), calcium channels (*TRPC6, ITPR1*) and sodium channels (*ASIC2*) (Fig. 4B). We also found increasing expression of a range of solute carrier (SLC) family genes, which use ion gradients to transport a range of cargo. Older ChP organoids (> day 70) showed increased expression of sodium symporters that transport iodine (*SLC5A5*), amino acids (*SLC6A15*), myo-inositol (*SLC5A3*), bicarbonate (*SLC4A5*), ascorbate (*SLC23A2*) and L-glutamate (*SLC1A3*), consistent with specialised barrier transport functions (Fig. 4B). This module also captured a temporal increase in the mitochondria fraction, reflecting the elevated metabolic capacity required for CSF secretion in mature ChP epithelium (Fig. S4B).

**Figure 4.**
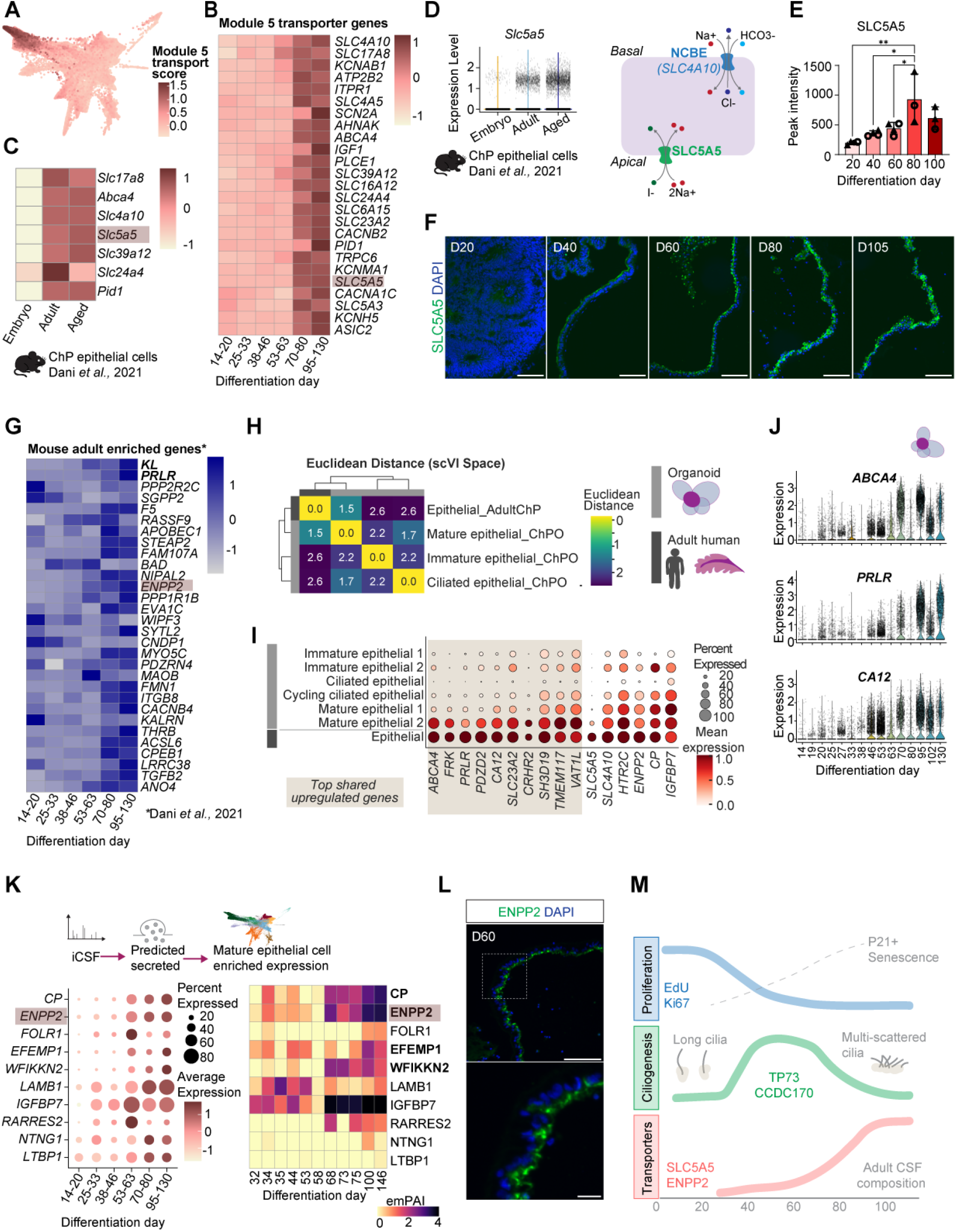
ChP organoids achieve mature transcriptional and functional identity. **(A)** FA plot coloured by geneset score for transmembrane transport genes (module 5). **(B)** Pseudobulk expression heatmap of epithelial module 5 transmembrane transport genes across time in ChP organoid epithelial lineage cells grouped by age. **(C)** Pseudobulk expression heatmap of module 5 transmembrane transport genes in mouse ChP epithelial cells across ages ^20^. Selected genes significantly increased in adult vs embryonic epithelial cells. **(D)** Violin plot of *Slc5a5* increasing with age in mouse ChP epithelial cells ^20^. Schematic indicates localisation of selected adult-enriched ion transporters on the ChP epithelium. **(E)** Quantification of SLC5A5 intensity measured by line profile perpendicular to the epithelial layer. Points represent average per organoid (n=3-4), cell lines are indicated by point shape: solid triangle H1, open circle H9. One-way ANOVA with Tukey HSD, * p <0.05 ** p< 0.01. **(F)** ChP organoid sections across differentiation immunostained for SLC5A5 (green). Scale bar: 100 µm. **(G)** Pseudobulk expression heatmap of mouse ChP epithelium adult-upregulated genes ^20^ in ChP organoids epithelial lineage over time, grouped by age. **(H)** Heatmap showing similarity between organoid mature epithelial cells and adult human ChP epithelial cells. Adult ChP single-nuclei RNA (snRNA) sequencing data Mount Sinai cohort. Colour scale represents Euclidean distance between cluster centroids in scVI latent space. **(I)** Dotplot showing gene expression of selected mature epithelial markers across organoid epithelial subtypes (light grey) and adult epithelial cells (dark grey). Adult ChP snRNA sequencing data Mount Sinai cohort. Top 10 shared upregulated genes in ‘Mature epithelial 2’-organoid cluster and ‘Epithelial’-adult cluster highlighted. Point size represents percentage of cells expressing gene and colour scale shows average expression. **(J)** Violin plots showing expression of selected markers of adult ChP epithelium across time in ChP organoid epithelial lineage cells. **(K)** Secreted CSF proteins with gene expression enriched in mature epithelial cells in ChP organoids. Left: Dotplot of gene expression in all cells over time in ChP organoids. Dot size indicates percentage of cells expressing the gene and colour scale shows average gene expression. Right: Heatmap coloured by protein levels detected in iCSF by mass spec ^63^. Proteins in bold were also detected in adult human CSF. **(L)** Day 60 ChP organoid immunostained for ENPP2 (green). Scale bar: 100µm, inset 20µm. **(M)** Schematic of changes to epithelial cells in ChP organoids over differentiation time. Nuclei labelled with dapi (blue).

To benchmark our late-stage organoid epithelium with *in vivo* mature ChP, we compared our atlas with a previously published mouse transcriptomic datasets ^20^. Of the module five transport related genes, 11 showed the same age-dependent expression in mouse ChP cells, with enrichment in adult and aged samples (Fig. 4C, Fig. S4A). We validated the progressive enrichment of one of these, SLC5A5, in ChP organoid at the protein level, observing increased expression over time (Fig. 4C-E). Across the broader transcriptional programme, genes enriched in adult versus embryonic mouse ChP epithelium increased progressively over organoid development and were specifically enriched in the mature ChP epithelial clusters (Fig. 4G). Among these genes we found an increase in Klotho (*KL*), a canonical anti-ageing factor secreted into CSF ^45,46^, *PRLR*, encoding the prolactin receptor, which is most highly expressed in the brain at the ChP and mediates prolactin transport from blood into CSF ^47^ and *ENPP2* (autotaxin), a CSF-secreted protein that plays a critical role in the production of lysophosphatidic acid required for postnatal oligodendrocyte differentiation and myelination ^48^ (Fig. 4G).

We next investigated whether organoid epithelial cells could reach human adult transcriptional identity. We integrated our ChP organoid data with an adult human ChP snRNAseq dataset (Mount Sinai cohort, Pjanic et al., 2026, submitted), considering dataset and sequencing platform effects. The large cohort study (48 donors) combined more control donors of a range of ages (median 67.5 yr, range 23-101) than previous studies, providing a representative human ChP atlas for comparison. From the UMAP cell clustering, we observed similarities between organoid mature epithelial cells and adult epithelial cells, and between organoid stromal cells and adult mesenchymal clusters (Fig. S4C-E). As expected, both datasets also contained unique cell types, such as immune and vasculature in the adult, and progenitor and hem populations in the organoid (Fig. S4C-D). Utilising more dimensions, we calculated the difference between cell types as the distance between centroids of each cluster in scVI latent space (Fig. S4E). A shorter distance, and hence greater similarity, was found between the adult human epithelial cells and the mature organoid epithelial cells consistent with the full differentiation of these cells (Fig. 4H, Fig. S4F). Organoid mature epithelial cells and adult epithelial cells shared high expression of genes functioning in CSF production *SLC23A2*, *PRLR*, *CA12* and genes involved in neuroendocrine sensing *HTR2C and CRHR2* (Fig. 4I, 4J).

We next asked whether this maturation was reflected in the organoid iCSF profile. We found that 83.7% of the iCSF proteins from late-stage organoids (d68-d146) is present in human adult CSF (Pellegrini et al 2020; Fig S4G). As expected, proteins present only in human adult CSF but absent in the organoid iCSF, revealed enrichment for blood/plasma components (Fig.S4H). Consistent with our transcriptomic data, we observed progressive enrichment in ENPP2 (Fig.4L) and ceruloplasmin (CP), a ferroxidase secreted by ChP epithelial cells into CSF that regulates brain iron homeostasis ^49^, further supporting the acquisition of a mature secretory function (Fig. 4K).

In summary, these data define an ordered developmental programme in which human ChP organoids sequentially achieve hallmarks of *in vivo* maturation: an early proliferative decline, coordinated ciliogenesis and progressive acquisition of transmembrane ion channels and transporters characteristic of the mature postnatal epithelium (Fig. 4M) ^1,14,20^.

### VPA reveals a stage-specific and reversible plasticity in human ChP maturation

Having defined the ordered phases of human ChP epithelial maturation, we next asked whether this programme retains the capacity to recover from perturbation applied during defined developmental windows. To test this, we used valproic acid (VPA), HDACs inhibitor ^50,51^ whose *in utero* exposure is associated with neurodevelopmental defects ^52^. VPA reversibly perturbs chromatin regulation ^53–55^, and it provides a tractable tool to test the sensitivity and recoverability of distinct maturation phases. Although the ChP has recently been identified as a specific target of gestational VPA in rat ^19^, and in human cortical organoids ^62^, whether human ChP maturation is affected by VPA and whether this is dependent on a specific stage of development has not been examined.

We applied VPA during two developmental windows: an early window (day40 onset) coinciding with multiciliogenesis, and a late window (day60 onset) coinciding with secretory maturation, each at short (20-day) and long (40-day) treatment durations (Fig. 5A). To test reversibility, we washed out VPA after 20 days of exposure in both early and late conditions and assessed the organoids following a 20-day recovery period.

**Figure 5.**
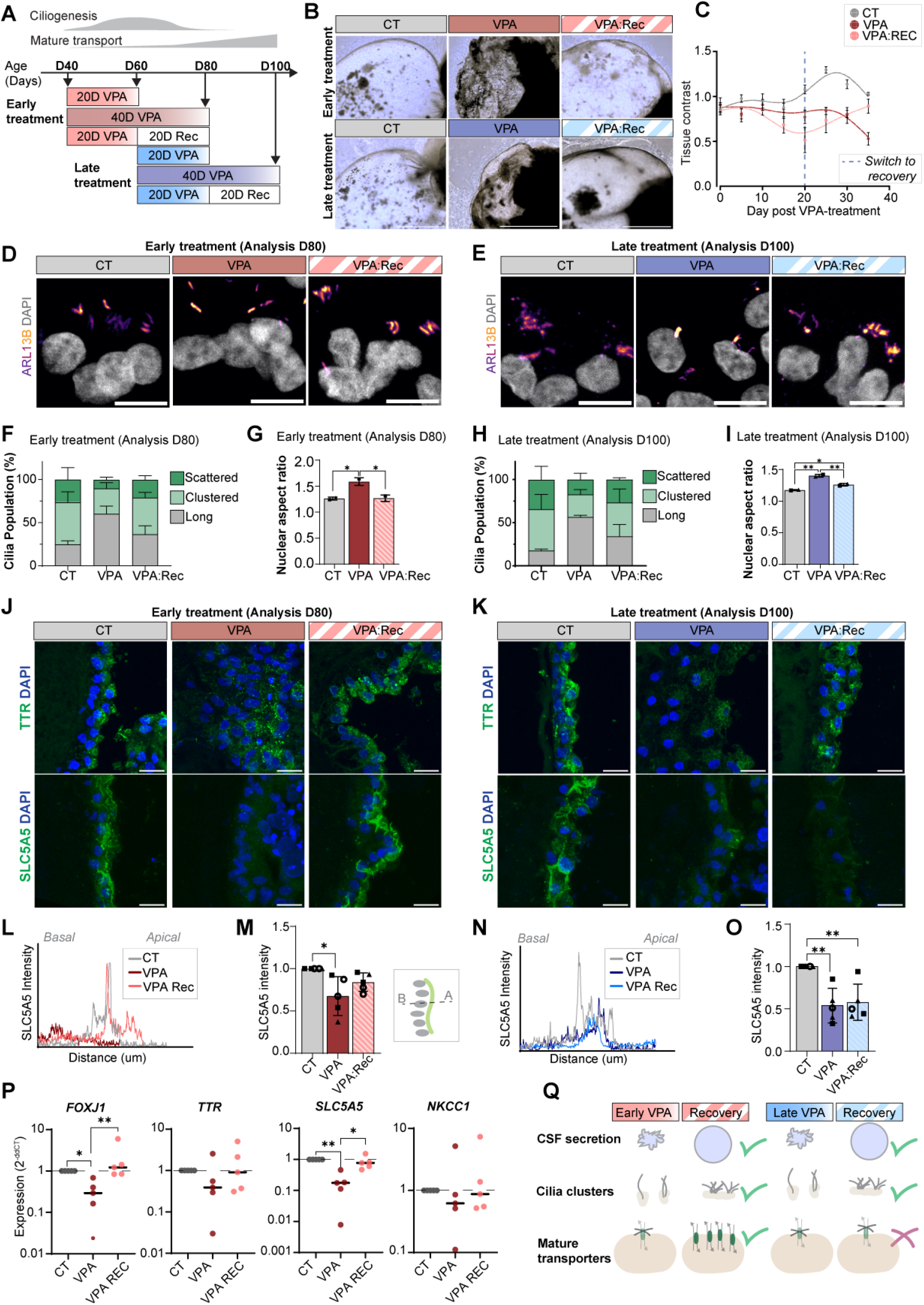
Valproic acid disrupts ChP cilia and secretion with developmental stage-dependent recovery. **(A)** Schematic of VPA treatment. ChP organoids were treated with 1mM VPA from D40 (Early Treatment) or D60 (Late Treatment). Treatment lasted 20 days before collection or a 20-day recovery (Rec). Alternatively, treatment lasted 40 days. **(B)** ChP cyst collapsed after VPA treatment at both time windows and recovered after VPA removal only after early treatment, representative brightfield images. **(C)** Cyst tissue contrast over time for early VPA treatment, starting at D40. Points represent mean value with standard error bars. Transition from VPA to recovery for VPA:Rec samples indicated by dashed line. **(D)** D80 ChP epithelial cilia (ARL13B, fire) across conditions of early VPA treatment. Nuclei stained with dapi, grey. Scale bar: 10µm. **(E)** D100 ChP epithelial cilia (ARL13B, fire) across conditions of late VPA treatment. Nuclei stained with dapi, grey. Scale bar: 10µm. **(F)** D80 ChP organoid cilia classified according to morphology described in Figure 3E. Proportion of cilia across treatment conditions, plotted as mean with standard error bars. Two-way ANOVA and Tukey multiple-comparison test, n=4 with 30-50 cells/condition/experiment. Long cilia significantly increased after 40 days of VPA exposure (adj. p < 0.0001 for VPA vs CT, p <0.01 for VPA vs VPA:Rec), while all cilia populations returned to control levels after recovery (adj. p = n.s for VPA:Rec vs CT). **(G)** Nuclear aspect ratio (longest axis / shortest axis) across conditions for early VPA treatment (n=2). **(H)** D100 ChP organoid cilia classified according to morphology described in Figure 3E: long cilia (grey) short, clustered cilia (light green) and scattered multi-cilia (dark green). Proportion of cilia across treatment conditions, plotted as mean with standard error bars. n=2 with 30-50 cells/condition/experiment. **(I)** Nuclear aspect ratio (longest axis / shortest axis) across conditions for late VPA treatment. Points show average for batch, n=2. **(J)** Early VPA exposure decreased expression of secretion markers (green) in D80 ChP epithelium and expression was restored in recovery condition. Immunostained against TTR (top) and SLC5A5 (bottom). Nuclei labelled with dapi (blue). Scale bar: 20µm. **(K)** Late VPA exposure decreased expression of secretion markers (green) in D100 ChP epithelium and remained low in recovery condition. Immunostained against TTR (top) and SLC5A5 (bottom). Nuclei labelled with dapi (blue). Scale bar: 20µm. **(L)** Apical localisation of SLC5A5 is disrupted by early VPA treatment, measured at D80. Example intensity profile of SLC5A5 along a line perpendicular to ChP epithelium. **(M)** Peak SLC5A5 intensity across early VPA treatment conditions at D80. Normalised to control (n=5). **(N)** Apical localisation of SLC5A5 is disrupted by late VPA treatment, measured at D100. Example intensity profile of SLC5A5 along a line perpendicular to ChP epithelium. **(O)** Peak SLC5A5 intensity across late VPA treatment conditions at D100. Normalised to control (n=5). **(P)** Expression of selected genes following early VPA treatment, measured at D80. Expression calculated as 2^-ddCT^, normalised by housekeepers and relative to control (n=5). **(Q)** Schematic of changes to ChP following VPA treatment and treatment washout at early and late timepoints. Data point on graphs represent independent organoid batches (“n”). Cell lines are indicated by point shape in G, I, M & O: solid triangle H1, open circle H9 and solid square KOLF2.1J. Statistical test used across three conditions: One-way ANOVA and Tukey multiple comparison test. *p <0.05, **p<0.01, ***p<0.001.

VPA treatment caused the organoid fluid-filled cysts to collapse at both treatment timepoints, with reduced cyst size and tissue contrast (Fig. 5B, C). Cyst collapse was reversed after VPA washout at both timepoints. Despite the morphological disruption, the ZO1 tight junction organisation remained intact across conditions, indicating that loss of cyst volume was not caused by barrier disruption (Fig. S5A).

Given previous reports of VPA effects on ciliogenesis ^19,56^, we next examined ciliary morphology in the ChP epithelium following VPA treatment and washout. Both early and late VPA exposure reduced multiciliated cilia populations (scattered and clustered), while increasing long cilia, an effect that was reversed after washout (Fig. 5D-I). VPA also increased nuclear elongation, opposing the maturation-associated transition to rounded nuclei, and this phenotype was similarly reversed after washout (Fig. 5G, I, and Fig. S5B-E). Consistent with these protein changes, transcript levels of the multiciliogenesis regulator, *FOXJ1* were similarly disrupted (Fig. 5P). These findings show that VPA transiently affects ciliogenesis and nuclear morphology, which may reflect VPA-mediated HDAC inhibition leading to microtubule-stabilisation. These cytoskeletal phenotypes recover following drug withdrawal irrespective of the ChP developmental window of exposure.

To determine whether the mature transporter programme showed similar reversibility to this perturbation, we examined the expression of CSF protein, TTR, and apical transporter, SLC5A5. VPA exposure reduced both markers during early and late treatment (Fig. S5F). In contrast to the reversible cyst and ciliary phenotypes, recovery of transporter expression was stage-dependent: TTR and SLC5A5 expression was restored back towards control levels following early VPA withdrawal, while both markers remained reduced in expression after late VPA washout (Fig. 5J-O). Transport-related genes *SLC5A5, TTR* and *NKCC1* were reduced at a transcript level after early VPA treatment (Fig. 5P), similarly indicating disruption of the epithelial secretory state.

Together, these findings reveal a stage-specific plasticity in human ChP maturation: ciliogenesis and nuclear morphology fully recover after transient exposure at any developmental stage, whereas the mature transporter programme re-engages only when disruption occurs before the secretory maturation is established (Fig. 5Q).

## Discussion

Despite its role in CSF secretion, blood-CSF-barrier and brain development, the maturation programme of the human ChP has remained largely inaccessible to mechanistic investigation. Here, we combine longitudinal scRNA transcriptomics of ChP organoids compared against human foetal and adult ChP, centrosome-enriched proteomics, and pharmacological perturbation to resolve this programme at unprecedented temporal resolution. Our data reveal that human ChP epithelial maturation proceeds through sequential phases from proliferation exit to multiciliogenesis to secretory maturation, culminating in a transcriptional state concordant with mature human ChP function. We further identify a stage-specific asymmetry in the reversibility of these phases on VPA treatment: ciliary and nuclear morphology disruptions remain reversible throughout the maturation window, while disruption of the mature secretory programme fails to recover once initiated. Together these findings define both the ordered architecture of human ChP maturation and the temporal boundary at which it becomes less plastic.

Prior characterisation of adult human ChP ^57^ and developing mouse ChP ^20^ is available, but a longitudinal human developmental atlas has not been previously shown. Our 15-timepoint dataset reveals that organoids recapitulate the remarkable proliferative decline, including the directional incorporation of new cells into mature epithelium from the root of the plexus. This rapid proliferation exit is accompanied by a p21-associated terminal differentiation signature recently described in the maturing mouse ChP ^1,15,24^. This suggests that p21-mediated cell cycle exit is a conserved developmental mechanism by which ChP epithelium commits to a stable post-mitotic state.

The acquisition of multiple nodal-like apical cilia which regress over time during development is a defining feature of ChP epithelial identity and is critical for CSF flow, mechanosensation and signalling ^1,58^. Our pseudotime analysis identifies a ciliogenesis gene module (CFAP73, FOXJ1, DNAI2, CCNO) peaking between days 40 and 60. Cilia reach similar length and spatial organisation comparable to what has previously been reported in postnatal ChP ^1^. Using CAPture proteomics, we identified novel centrosome associated proteins, including MYH14 and CCDC170, with no prior characterisation in the ChP, but with stage-specific temporal centrosome recruitment patterns. These findings provide the first available dataset of the centrosomal proteome across development in human ChP.

A central limitation of current organoid systems has been their predominant alignment with foetal rather than adult tissue transcriptional states ^59^. Our data demonstrate that, within this model, some ChP epithelial cells reach a transcriptional state that aligns with adult human ChP tissue representing the first stem cell derived ChP model to achieve a postnatal-like maturation endpoint. This is supported by several lines of evidence: scRNAseq data, proliferation analysis using reporter lines and EdU incorporation, and late-stages organoid epithelium showing progressive enrichment of transporters characteristic of mature ChP *in vivo* (using both human and mouse datasets), confirmed at both transcriptional and protein levels. The shared functional characteristics with adult tissue further establishes the late-stage organoid system as a relevant *in vitro* platform for studying mature human ChP biology, including processes that have until now been inaccessible to mechanistic investigation in human systems.

To probe the developmental architecture of this maturation programme, we used VPA as a pharmacological perturbation. VPA effects are reversible upon washout in hepatocytes and in adult cortical tissue, where VPA-induced chromatin opening transiently reactivates critical period plasticity ^55,60^. VPA is also widely used as a reversible chromatin-modifying agent in iPSC reprogramming ^61^. This pharmacokinetic reversibility, combined with VPA’s chromatin mechanism, makes it well suited to test whether developmental programmes that have been transiently disrupted can re-engage upon drug withdrawal. We find that VPA disrupts both multiciliogenesis and secretory maturation during exposure, but we observed different recovery phenotypes: ciliation and nuclear elongation recover fully after washout at all developmental stages, while recovery of mature transporter SLC5A5 is limited to the early treatment window. Consistent with our findings, recent studies have shown that gestational VPA exposure induces transcriptomic changes in the neonatal rat ChP ^19^ and VPA-induced transcriptional changes have been identified in cortical organoids, including in a small ChP-like cluster ^62^. The reversible switch from multiciliated to monociliated cells, together with the nuclei elongation, support the homeostatic plasticity of these processes. However, the failure to recover SLC5A5, within the experimental window, indicates that not all VPA effects are equally reversible.

This asymmetry of recovery of ciliogenesis and transporter modules, suggests that these two phases differ not only in their developmental timing, but also in their regulatory architecture. These findings suggest that ciliogenesis is an ongoing structural process the cells can re-execute throughout maturation. Whilst secretory maturation represents a single developmental transition that fails to re-establish once the developmental window is closed.

Our work provides a systematic developmental framework for human ChP biology and has direct implications for understanding when and how teratogen-induced ChP dysfunctions become irrecoverable. It also establishes a recoverability paradigm using timed perturbation and washout to distinguish reversible from durable developmental disruptions in an experimentally tractable human organoid system, opening this approach to broader applications across other tissues and other teratogens or environmental stressors.

## Limitations of the Study

There are several limitations that warrant consideration. First, our organoid system models human ChP epithelial development but they do not recapitulate full physiological context and lack immune surveillance, CSF circulation and vascular interactions. Second, while our data show that disruption of the mature transporter SLC5A5 does not recover during the experimental window assessed, we cannot fully exclude the possibility of recovery on longer timescales. Testing this would require extended culture experiments beyond practical limits of this study. Third, although the asymmetric reversibility we observe is consistent with chromatin-level regulation, we cannot definitively attribute VPA effects to HDAC inhibition nor we directly demonstrate the underlying chromatin regulation, and future work using ATAC-seq, ChIP-seq or specific HDAC inhibitors would be warranted to mechanistically address this. Fourth, while we benchmarked ChP developmental maturation against both foetal and adult human tissue, direct *in vivo* validation of VPA-induced phenotypes, was not feasible due to limited access to human tissue. Fifth, the centrosomal proteins we identified are presented as candidates which will require further validation through knockout or perturbation approaches that are beyond the scope of this study. Efforts to validate additional CAPture candidates were limited by the availability of specific antibodies that reliably recognised the target proteins. Finally, direct benchmarking against primary human tissue remains constrained by the limited availability of ChP samples, particularly across fetal and early postnatal development. Expanding these reference datasets across developmental stages is the focus of ongoing work in our laboratory. Despite these limitations, our findings establish the foundation for understanding the developmental architecture of the human ChP and provide a framework for investigating its regulatory boundaries under developmental perturbations.

## Resource availability

## Lead Contact

Requests for further information and resources should be directed to and will be fulfilled by the lead contact, Laura Pellegrini.

## Materials availability

This study did not generate new unique reagents.

## Data and code availability (GEO submission)

GEO accession numbers: GSE330823 and GSE330822 (10X and Parse) and previously published (GSE150903). Code of the single cell analysis is available on GitHub (https://github.com/LizzyApsley/ChP-organoid-timecourse).

## Acknowledgments

The authors would like to thank members of the Pellegrini Lab for the helpful feedback and discussions. This study was supported by Wellcome Trust funding (225950/Z/22/Z). K.R.L. would like to thank Andrew Copp, Deborah Henderson, Steve Lisgo, Nita Solanky and their teams at the HDBR for their invaluable support, and the funding sources the Medical Research Council (MRC) MR/S025065/1 and Heart of Racing. FG and EV are supported by the MRC (UKRI585). MAL is supported by the MRC (MC_UP_1201/9) which provided funding for the single-cell RNA sequencing performed in this study.

## Author contributions

E.A. designed and conducted experiments, analysed data and wrote the manuscript. S.S.T. designed and conducted VPA experiments, cilia imaging and data analysis. I.I.R. conducted initial bioinformatic analysis and data integration. E.V. conducted CAPture proteomic workflow and analysis. K.S. prepared and maintained organoid cultures and provided technical support. M.F.O. prepared and maintained organoid cultures, processed human tissue samples and provided technical support. M.P., J.B., J.F.F., and P.R. contributed the Mount Sinai cohort data, including data generation, processing, and curation, and participated in manuscript review. F.G. supervised CAPture work and participated in manuscript review. K.R.L. provided human tissue samples and participated in manuscript review. M.A.L. helped with collection of the scRNA-seq samples and participated in manuscript review. L.P. designed and supervised the project, analysed data and wrote the manuscript.

## STAR Methods

### EXPERIMENTAL MODEL AND STUDY PARTICIPANT DETAILS

Human embryonic stem (hES) cells H9 and H1 (WiCell WA09, WA01) and the human induced pluripotent stem cell (hiPSC) line KOLF2.1J (Jackson Laboratory, JIPSC1000) were approved for use by the Medicines and Healthcare products Regulatory Agency (Reference #SCSC23-57). Cell lines are indicated by point shape in plots: solid triangle H1, open circle H9 and solid square KOLF2.1J. H9 FUCCI reporter line was generated in the Lancaster lab and shared under MTA (Reference #OG5995). Cells were cultured in 6-well plates coated in Matrigel Growth Factor reduced (GFR) Basement Membrane Matrix (Corning, 356230) and maintained in StemFlex medium (Gibco, A3349401). Embryoid bodies (EBs) were prepared from single cell suspension from hES as previously described ^1^. STEMdiff Choroid Plexus Organoid Differentiation Kit (STEMCELL technologies, 100-0824) reagents were used for the generation of ChP organoids following kit instructions. Following ChP patterning on day 10, EBs were kept in a 5cm dish and then moved to the shaker on day 14 to be fed every 3-4 days with maturation media. From day 30 dissolved Matrigel (1:50-1:100, Corning, 354234) was added to the maturation media.

### Human foetal choroid plexus tissue

Human foetal tissues were obtained from the Human Development Biology Resource (HDBR), provided by the Joint MRC/Wellcome Trust (grant #MR/R006237/1) (www.hdbr.org) in collaboration with Dr Katherine Long’s lab. The HDBR provided fresh tissue from foetuses aged 8–14 post-conception weeks. All tissue was fixed for at least 24h at 4°C in 4% (wt/vol) paraformaldehyde (PFA) in 120 mM phosphate buffer (pH 7.4).

## METHOD DETAILS

### Immunostaining

Organoids were fixed in 4% PFA overnight at 4°C or for 1 h at room temperature then washed with PBS and moved to 30% sucrose buffer in PB at 4°C for at least 24 h. The following procedure was used for organoid and human foetal samples. Organoids or human foetal tissue were embedded in a warm solution with 7.5% gelatin 10% sucrose in PB at 37°C and frozen blocks were sectioned as previously described ^1,2^. Slides were thawed at room temperature, washed with PBS and incubated with permeabilization/blocking solution (4% NDS, 0.25% Triton-X-100, PBS) at room temperature for 1 hour. Next, primary antibodies were diluted in antibody solution (4% NDS, 0.1% Triton-X-100, PBS) overnight at 4°C. After washing three times with PBS, secondary antibodies were applied 1:1000 in antibody solution for 1h at room temperature. Slides were washed in PBS again three times, with DAPI added 1:1000 in the last wash, before mounting coverslips with a mounting medium (Prolong Diamond).

### Confocal imaging

Images were taken using a LSM 800 Axio imager (Zeiss), Eclipse Ti-2 inverted confocal microscope with SoRa super resolution disk (Nikon) or AXR Ti-E 2 inverted confocal microscope (Nikon). ImageJ/Fiji (v1.53f51) was used for image analysis.

### Single cell RNA sequencing

Single cell dissociation was performed as previously described ^3^ (Table S1). Briefly, samples were incubated in 1ml of Accumax (Sigma, A7089) with 400μg DNase I and 15μM actinomycin D at 37°C for 20min with gentle agitation and intermittent pipetting every 5 min. 100μl FBS was then added to the cell suspension before filtration on a 35μm filter tube (Corning, 352235).

Samples were then spun at 300g for 5min. For 10X Genomics analysis, cells were resuspended in an appropriate volume of 0.04% BSA in PBS to load 16,000 cells per well on the 10X Chromium system (10X Genomics). 10X sequencing was performed at CRUK Cambridge by the Genomics core facility. Single cell RNAseq libraries were prepared according to manufacturer’s instructions using the 10X Genomics Chromium Single Cell 3’ Library & Gel Bead Kit v3 (10X Genomics) workflow. The Chromium Controller produced the single cell beads. Reverse transcriptase reaction and subsequent amplification was carried out on a C1000 Touch Thermal Cycler (Biorad). Libraries were quality tested using a 2100 Bioanalyzer Instrument (Agilent). Samples were sequenced in two rounds on one lane each of the SP flowcell of the Novaseq6000 sequencer (Illumina).

For Parse Bioscience analysis, cell suspension was spun at 200g for 10min at 4°C and the pellet resuspended in cell prefixation buffer (Parse Bioscience). Sample fixation and split-seq barcoding were performed according to the Evercode Fixation v2 Parse Bioscience protocol. Single cells were sequenced with S2 Novaseq lane (4 billion read depth). Single nuclei were sequenced with SP Novaseq lane (400 million read depth).

### Bioinformatic analysis

#### Processing of Parse Bioscience scRNAseq

Raw sequencing data from Parse Biosciences was processed using Parse Biosciences software split-pipe v0.9.6p. Alignment was performed against the human genome reference GRCh38 and genome annotation GRCh38.93. A filtered digital expression matrix (a cell-by-gene count matrix) was created using combinatorial barcoding containing UMI counts with split-pipe default QC parameters. Cell barcodes with less than 500 genes, mitochondrial fraction below 20% were excluded. Transcriptome size factors were calculated using computeSumFactors from the R scran package ^4^ Cells were pre-clustered with scran’s quickCluster function (using the igraph method), with minimum and maximum cluster sizes of 100 and 3,000 cells, respectively. The resulting log2 normalised expression values were used for dimensionality reduction in the clustering step during doublet detection. Doublet calling was performed using the function scDblFinder ^5^, with fixed expected doublet rate of 3%.

#### Processing of 10X Genomics scRNAseq

Raw sequencing data from 10X Genomics publicly available (GSE150903) and newly generated was processed using Velocyto 10X UMI count software against CellRanger genome build GRCh38-1.2.0, including the GRCh38 repeats masker file ^6^. The resulting spliced and unspliced count matrices were aggregated for further analysis. Cell barcodes with less than 500 genes, mitochondrial fraction below 20% were excluded. Doublet calling was performed using the function scDblFinder ^5^ with default parameters. Approximately 9% of cells in the data were labelled as doublets.

#### Integration of Parse Bioscience and 10X Genomics data

UMI count data from Parse Bioscience and 10X Genomics was combined by shared gene ids. Integration was performed using scVI (via scvi-tools version 1.0.3) and Scanpy (version 1.9.4) to generate a shared, low-dimensional latent space corrected for two batch covariates (Individual datasets and sequencing platforms) while preserving biological variation and model gene expression (denoised counts). For dimensionality reduction and visualization the top 2000 highly variable genes were selected using Seurat_v3 method as implemented in Scanpy, followed by scVI integration. The resulting latent space was used for clustering with Leiden algorithm ^7^ as well as for generating the UMAP (with min_dist=0.3) and Force directed (FA) layouts used in the manuscript figures.

#### Cell type annotation and plotting

Leiden clusters were manually annotated using cluster markers and known cell type markers from literature. Leiden cluster 17 was excluded from subsequent analysis due to a high doublet score relative to all other clusters. Leiden clusters with consistent markers within cell types of interest were merged for the final broad cell type annotation. A Seurat object was generated with combined count data and log normalised data was used for plotting gene expression (Seurat version 5.1.0). Seurat, ggplot2 (version 3.5.1) and pheatmap (version 1.0.12) were used to generate figure plots.

#### Label transfer from mouse embryonic scRNAseq

Embryonic mouse ChP single-cell RNA-sequencing data ^8^ was filtered for lateral ventricle samples. Counts from mouse and organoid cells were combined using shared genes. Following batch correction, label transfer from mouse to organoid cells was performed using Seurat (Seurat_5.1.0) functions FindTransferAnchors and TransferData. A threshold prediction score of 0.5 was applied for label transfer.

#### Integration with 2D ChP differentiation protocol

Organoid data was integrated with single-cell RNA-sequencing data from a 2D ChP differentiation protocol ^9^. Counts from organoid and 2D data were combined as a Seurat object (Seurat version 5.1.0) using shared genes and split into three: organoid-10X, organoid-ParseBio and 2D-10X. Variable genes were identified for each dataset and SelectIntegrationFeatures() was used to select 2000 common highly ranked variable genes. The common highly variable genes were used to find integration anchors (reduction method ‘rpca’). Integration was performed sequentially, first joining the organoid platforms and then the 2D dataset. The integrated assay was used for dimensionality reduction and correlation of cell types. To compare cell types, cluster-level pseudobulk expression was generated using the integrated assay data and Pearson correlation calculated between clusters. Expression of example genes was plotted using the RNA assay log normalised data.

#### Epithelial lineage trajectory

Epithelial lineage cells were defined as hem and epithelial clusters, and subset from the full organoid dataset. A trajectory was fitted through the middle of the epithelial lineage in FA embedding coordinates using principal_curve (princurve version 2.1.6). Pseudotime scores for individual cells were calculated as distance along the trajectory normalised between 0 and 1. A CellDataSet object (monocle version 2.32.0) was constructed containing expression counts for the top 2000 variable genes by epithelial lineage cells, as identified using Seurat. Genes differentially expressed over pseudotime were identified using Monocle’s differentialGeneTest comparing a model dependent on pseudotime coordinates with a reduced model of constant expression. For the top 500 genes differentially expressed over pseudotime, smoothed expression curves over 200 pseudotime points were generated using genSmoothCurves. These genes were then grouped into five modules based on expression over pseudotime using cluster::pam (cluster version 2.1.6) function with a Pearson correlation based distance metric (Table S3). k = 5 was selected to capture the major patterns observed in expression over pseudotime. Gene ontology analysis was performed using gprofiler2 (version 0.2.3) for each module, using the Benjamini-Hochberg/FDR method for multiple-testing correction, with the 2000 variable genes as the background list and pathways from GO molecular function and GO biological processes (Table S4). Module scores were calculated using AddModuleScore from Seurat.

#### Plotting of mouse ChP snRNAseq data

Mouse ChP single-nuclei RNA-sequencing data ^10^ including three ages: embryonic, adult and aged was downloaded from https://singlecell.broadinstitute.org/single_cell/study/SCP1366/choroid-plexus-nucleus-atlas.

#### Adult human data integration

Adult human ChP data was obtained from a large disease cohort study (Pjanic et al., 2026, Manuscript submitted for publication at Cell). Specimens were sourced from Mount Sinai NIH Neurobiobank. From the full cohort, 48 donors (21 female, 27 male) were classified as control by three dementia scores (Braak neurofibrillary tangle stage, Braak; CERAD neuritic plaque score, CERAD; and Clinical Dementia Rating, CDR). These control donors were subset from the full processed dataset and used for comparison with organoid (Table S6).

Adult human snRNAseq library was prepared as follows. All buffers were supplemented with RNase inhibitors (Takara). 6 samples were processed in parallel. ∼25 mg of frozen tissue from each specimen was homogenized in cold lysis buffer (0.32 M Sucrose, 5 mM CaCl_2_, 3 mM Magnesium acetate, 0.1 mM, EDTA, 10 mM Tris-HCl, pH8, 1 mM DTT, 0.1% Triton X-100). After filtering through a 40 μm cell strainer, the flow-through was underlaid with sucrose solution (1.8 M Sucrose, 3 mM Magnesium acetate, 1 mM DTT, 10 mM Tris-HCl, pH8) and centrifuged at 107,000 × g for 1 hour at 4 °C. Pellets were resuspended in PBS and quantified using a Countess II cell counter (Life Technologies). 2 M nuclei from each sample were then pelleted at 500 × g for 10 minutes at 4 °C and re-suspended in 100 μl staining buffer (2% BSA, 0.02% Tween-20, 10 mM Tris, 146 mM NaCl, 1 mM CaCl_2_ and 21 mM MgCl). Each sample was incubated with 1 μg of a distinct TotalSeq-A nuclear hashing antibody (Biolegend) for 30 min at 4 °C. 7-AAD (Invitrogen) was added to each sample and 7-AAD positive nuclei sorted into tubes pre-coated with 5% BSA using a FACSAria flow cytometer (BD Biosciences). Following FANS, nuclei were washed in a staining buffer, pelleted, re-suspended in PBS and quantified. Nuclei concentrations were normalized and equal amounts from each specimen pooled together. 2 aliquots of 60,000 pooled nuclei (i.e. 10,000 per sample) were processed in parallel using 3′ v3.1 reagents (10x Genomics) to create technical replicates. At the cDNA amplification step (step 2.2), reactions were supplemented with a hash-tag oligo (HTO) cDNA “additive” primer (GTGACTGGAGTTCAGACGTGTGCTCTTCCGAT*C*T; *Phosphorothioate bond). Following cDNA amplification, supernatants from the 0.6x SPRI selection step were retained for HTO library generation. Otherwise, cDNA libraries were prepared according to the manufacturer’s instructions (10x Genomics Chromium v3). HTO libraries were prepared as described previously ^11^.

Sequencing reads were aligned to hg38 reference genome using STARsolo. Nuclei were assigned to their origin donor using a genotype-informed demultiplexing strategy followed by validation via genotype concordance with HTO labels. A standard Scanpy workflow was followed to merge donor files into a unified dataset. Cell and gene filtering thresholds were then applied to the full dataset: cells expressing fewer than 1000 genes or more than 10,000 genes were excluded; cells with a mitochondrial read fraction exceeding 10% excluded; genes detected in fewer than 3 cells were removed. Doublet detection was performed using Scrublet and per-cell QC metrics calculated using scanpy.pp.calculate_qc_metrics(). A shared nearest-neighbour graph was constructed in the Harmony-corrected PCA space and used for UMAP projection and Leiden clustering (resolution 0.1) to assign nuclei to broad cell types.

Adult and organoid counts for shared genes were combined and datasets integrated in python using Scanpy (version 1.11.5) and scVI (version 1.4.2). Variable genes were found independently for each dataset using Scanpy ‘Seurat_v3’ method and the intersection of 2399 highly variable genes used for integration. The scVI model corrected for origin (adult, organoid) as batch variable and dataset sequencing platforms as a covariate (10X organoid, Parse organoid, 10X adult). The resulting scVI latent space was used to generate a UMAP layout, calculate Euclidean distances between centroids of each cell type and generate denoised expression. Euclidean distances were calculated using cdist from scipy (1.17.1) and clustering hierachy based on linkage method ‘ward’. Plots were generated using matplotlib (version 3.10.8) and seaborn (version 0.13.2). To identify shared highly expressed genes between organoid ‘Mature epithelial 2’ and adult ‘Epithelial’ cells, denoised expression was used to calculate average expression for each celltype. Gene expression was then z-score transformed to highlight cluster-specific expression. Genes with highest values for z-score-’Mature epithelial 2-ChPO’ x z-score-’Epithelial-AdultChP’ were selected as upregulated in both cell types compared to the rest of the population. Packages anndata (version 0.12.10), numpy (version 2.4.3) and pandas (version 2.3.3) were used for data storage and manipulation.

#### Mass spectrometry CSF preparation and data analysis

Organoid CSF was collected using either a 1ml syringe or a pulled glass microcapillary attached to filter and tubing using controlled suction as previously described ^12^. Day 140 organoid iCSF was collected and human adult CSF (54y, male) was obtained from the London Neurodegenerative Diseases Brain Bank. CSF samples were prepared by in-solution reduction (DTT), alkylation (IAA), and overnight tryptic digestion (1:100 enzyme:protein ratio, 37°C) following acetone precipitation. Peptides were resuspended in 2% ACN/0.05% formic acid and separated by reversed-phase chromatography on a 75 µm × 50 cm C18 PepMap column (U3000 UHPLC NanoLC, ThermoFisher Scientific) over a 60 min acetonitrile gradient at 250 nl/min. Data-dependent MS/MS acquisition was performed on an Orbitrap Eclipse Tribrid (ThermoFisher Scientific) with full MS scans at 120,000 resolution (*m/z* 375–1800) and CID fragmentation at 35% collision energy. Raw data were searched against the UniProt Human database (51,624 entries) using SEQUEST in Proteome Discoverer v2.5 at 1% FDR, with carbamidomethylation (static), methionine oxidation, and N-terminal acetylation (variable) as modifications. Label-free quantification was used to assess relative protein abundance across samples. Protein identifications were compiled and integrated in a curated table with previously published proteomic resources for human adult CSF and for mature organoid CSF (from Pellegrini et al 2020, d68-146, Table S5). Gene symbols were used as the matching identifier, yielding 603 unique symbols for human adult CSF and 387 for mature organoid CSF.

#### EdU labelling

EdU labelling and detection was performed using the Click-iT EdU Imaging kit (Invitrogen, C10340) and protocol. For short term labelling, organoids were treated with 10μM EdU in medium for 24h, before changing to fresh medium. Samples were then collected and fixed after a further 24h (a total of 48 h after first EdU addition). For long term labelling, organoids were treated with 1μM EdU in medium for 1 week before fixing in PFA. All samples were processed for immunostaining as above. After secondary antibody staining, slides were post-fixed in 4% PFA for 20min at room temperature. Slides were then washed with PBS, incubated with 0.1 M glycine for 15min at room temperature and washed three times for 5min with 3% BSA/PBS. Next slides were incubated with the Click-iT reaction mix (1x reaction buffer, 4% CuSO_4_, 0.08% Alexa fluor azide, 1x buffer additive) for 30min at room temperature. Slides were washed three times with 3% BSA/PBS, incubated with dapi 1:1000 in PBS for 5min and washed in PBS before mounting.

#### Flow cytometry

Organoids were collected on day 7 (8-10 organoids), 14 (4-6 organoids), 35, 53 and 80 (2 organoids). After washing in PBS, organoids were incubated in 1 ml AccuMax (Sigma, A7089) at 37°C for 20min. Resuspension and gentle pipetting were used to aid dissociation. 100μl iDM+A medium was added and pipetting was used to dissociate to a single cell suspension. For hESCs, cells were washed in 0.5mM EDTA, then dissociated using 0.5ml Accutase for 3-4min before inactivating with 0.5ml StemFlex. The cell suspension was filtered through a 70 μM strainer and then spun at 200g for 4min. Cells were resuspended in FACS buffer (1% BSA (Sigma, A7906-100G) in PBS) at 0.5-1 x10^7^ cells/ml. To stain dead cells, Zombie UV™ Fixable Viability kit (Biolegend, 423107) was added 1:500 to 100μl of cell suspension and incubated for 15min in the dark at room temperature. Next 500μl of 3% BSA/PBS was added, before spinning at 300g for 4min and resuspending cells in 2% PFA. After 10min incubation in PFA, samples were spun at 300g for 5min and resuspended in 100μl of FACS buffer. Samples were stored in the dark at 4°C before analysis. Unstained and non-fluorescent cells (H1 hESC line) were used as negative controls and heat-treated cells as a positive control to set the live-dead gating. Analysis was performed on CytoFLEX LX analyser (Beckman Coulter).

#### Cilia CAPture experiment

For cilia CAPture analysis, multiple organoids were pooled at each timepoint from 5 independent batches, from two cell lines (H1, H9): 20 organoids at day 20 and day 40, 6-10 organoids at day 80. All organoids were processed for pulldown with the CAPture peptide or bead-only, followed by mass-spectrometry on a Bruker timsTOF Ultra2. One bead-only experiment was run for day 20, and 3 for day 40 and 1 for day 80.

Centrosomes were isolated using the CAPture method, previously described ^13^. Briefly, streptavidin-coated magnetic beads (Dynabeads™ M-280 Streptavidin; Invitrogen) were coupled to a biotinylated CCDC61-derived peptide (Biotin-SPSPTGGRALRFDPTAFVKAKERKQREIQMKQQ; >95% purity; Biomatik). For each pulldown, 60μl of magnetic beads were washed three times in TBS-N buffer (0.1% NP-40 in TBS) followed by a single wash in CAPture buffer (50 mM Tris-HCl pH 8.0, 300 mM NaCl, 0.2% NP-40, 10% glycerol, protease inhibitor cocktail (Complete EDTA-free, Roche), and phosphatase inhibitor cocktail (PhosSTOP, Roche)). Beads were subsequently incubated in CAPture buffer with 6 μl peptide (1 mg/ml stock) for 1.5h at 4°C with gentle rotation (12rpm), before washing four times in CAPture buffer.

Cells (3 x10^7^ per pulldown) were lysed in CAPture buffer on ice for 30min. Lysates were briefly sonicated and clarified by centrifugation (1800g for 10min at 4°C). Cleared lysates were then incubated for 2h at 4°C with either peptide-coupled beads or control beads lacking peptide, using end-over-end rotation. Following binding, beads were washed four times in CAPture buffer.

#### Cilia CAPture mass spectrometry

Beads were washed twice in 100mM ammonium bicarbonate and then subjected to on-bead proteolytic digestion with 100ng trypsin in 50mM ammonium bicarbonate (AMBIC; Sigma) overnight at 37°C. An additional aliquot of trypsin was subsequently added, and digestion was continued for a further 4h at 37°C. Following digestion, the peptide-containing supernatant was collected, and proteolysis was quenched by addition of formic acid to a final concentration of 0.5% (v/v). Peptides were desalted using in-house prepared C18 solid-phase extraction cartridges. Cartridges were activated with 100% acetonitrile and equilibrated with 0.1% formic acid prior to sample loading. After loading, the cartridges were washed with 0.1% formic acid, and peptides were eluted using 50% acetonitrile containing 0.1% formic acid. Eluted peptides were dried and subsequently submitted for MS analyses.

Peptide samples were analysed using a NanoElute 2 UHPLC coupled to a timsTOF Ultra 2 mass spectrometer using a CaptiveSpray ionisation source (Bruker). Peptides were separated on a 75μm × 25cm IonOptiks Aurora Ultimate C18 analytical column (IonOptiks). Buffer A was 0.1% FA in H2O and buffer B was 0.1% FA in LC-MS grade acetonitrile (MeCN). A 30 or 120-minute linear gradient (0% to 40% buffer B) was used for elution. Source parameters were set to a capillary voltage of 1,600 V, a dry gas flow of 3.0 L/min, and a dry gas temperature of 200 °C. Data was acquired in dia-PASEF mode over an m/z range of 400–1,000 and an ion mobility range of 1/K₀ = 0.64–1.45 Vs/cm2. The TIMS analyzer was operated at a 100% duty cycle with 100ms accumulation and 100ms ramp times (acquisition cycle time = 0.96s). Collision energies were linearly stepped from 20 to 59 eV as a function of increasing ion mobility.

DIA data were quantified using DIA-NN (v2.2.0) ^14^ with a spectral library predicted from FASTA (Homo sapiens, swiss-prot, 20431 protein entries). Mass accuracy settings for MS1 and MS2 were fixed at 15. Precursor and protein group identifications were filtered at 1% FDR. Match between runs and protein inference were enabled. QuantUMS (precision) was used for quantification with retention time dependent cross run normalisation. Quantification matrices for precursors, protein groups, and gene groups were exported with 1% FDR filtering. N-terminal methionine excision and carbamidomethylation of cysteines were set as fixed modifications.

#### Cilia CAPture analysis and data processing

Mass spectrometry protein MaxLFQ intensity values were analysed in R (v4.4.2). Data were filtered, retaining only proteins for downstream analysis if they were detected in all 3 replicates in at least one condition. As centriole number per cell increases substantially during differentiation into multiciliated cells, raw intensities were normalised to CEP135, a centriolar marker, to account for this centriole amplification and allow meaningful comparison of relative protein abundance across timepoints. Intensity values were log2-transformed and median-centred. Missing values were imputed randomly from a normal distribution downshifted from the observed condition mean by 2.32 standard deviations with a width of 0.3 times the standard deviation. A fixed random seed was used to ensure reproducibility. The average fold change was then calculated by comparing matched peptide vs bead-only control samples from each organoid harvesting time point. Only proteins with a 2.3x fold enrichment (log2 scale) in peptide over bead-only were taken forward for Venn diagram comparison.

To compare H1 and H9 d40 organoids, proteomic intensity data were analysed in R using a custom pipeline including preprocessing, missing-value imputation, and differential abundance analysis. Proteins detected in at all three replicates per group were retained, log2-transformed, median-centred, and missing values were imputed using a Perseus-style downshifted normal distribution ^15^. Differential abundance testing was performed with the limma package using linear modelling and empirical Bayes moderation, with P values adjusted by the Benjamini–Hochberg FDR method. Proteins were considered significantly differentially abundant when adjusted P < 0.05 and absolute log2 fold change > 1.32.

#### VPA Treatment

To model the physiological exposure of valproate (VPA) on developing choroid plexus, VPA (P4543-25G, Millipore-Sigma) was added to the culture medium at a clinically relevant concentration of 1.0mM ^16^, with two media changes per week for the duration of the treatment. Organoids were treated with VPA from d40 (early treatment) or d60 (late treatment). A subset of the treated organoids was subsequently removed from VPA treatment (VPA washout) to study the effect of ChP recovery. Treatment were either stopped and samples were collected for post-processing at d80 or d100, depending on the initial exposure. Brightfield images were acquired on an EVOS XL Core microscope (Thermo Fisher, AMEX1000).

## QUANTIFICATION AND STATISTICAL ANALYSIS

### EdU frequency in epithelial cyst

Images were blinded before processing. Confocal z-stacks images were maximally projected. The epithelial cyst region was manually selected, viewing dapi and NKCC1 channels, and the ROI saved. DAPI ROI were selected using plugin StarDist 2D (model: Versatile (fluorescent nuclei), Percentile low: 1.0, Percentile high: 99.8, Probability Score/threshold: 0.5, Overlap threshold: 0.2) and filtered to those contained in cyst ROI. EdU ROI were provisionally selected using StarDist as above and then manually corrected if needed, before filtering to those contained in cyst ROI. Percentage EdU (Total EdU area / total dapi area x 100) was calculated per image, 3-5 images were averaged per sample.

### EdU distribution across cyst

Confocal z-stack images were maximally projected. Using the dapi and TTR channels, a freehand line was drawn along the complete length of the cyst and coordinates saved. Next, EdU+ cells along the cyst were selected using the multipoint tool and coordinates saved. After loading in R, coordinates were formatted as a spatial line (linestring object) and a list of points in the same geometric space. EdU points were snapped to their nearest points on the reference line using sf::st_nearest_points(). The relative distance (d) of each mapped point along the reference line was calculated using sf::st_line_project(). The relative distance from the nearest end (D) was then calculated for each point: if d < 0.5 D = d, else if d > 0.5 D = 1-d. A mean relative distance from near end was then calculated using all EdU points per image. EdU distances were averaged from 2-4 images per organoid.

### p21-positive nuclei

Confocal z-stack images were maximally projected and epithelial ROIs selected. Nuclei ROI were selected using plugin StarDist 2D (model: Versatile (fluorescent nuclei), Percentile low: 1.0, Percentile high: 99.8, Probability Score/threshold: 0.5, Overlap threshold: 0.6) and filtered to epithelial regions. p21 intensity was measured in all ROI and scored as positive above threshold value set using Otsu.

### TP73 intensity

Confocal z-stack images were maximally projected. To select nuclei, a threshold (method = Otsu) was applied to the dapi channel, followed by analyse particles (min size = 20). Nuclei ROI were checked and corrected manually if necessary, including limiting to the epithelial layer. In the TP73 channel, raw integrated density and area was measured for nuclei ROI and used to calculate mean TP73 fluorescence intensity (raw integrated density/area).

### Cilia length and classification

For cilia imaging, images were acquired at 12-bit, 512×512 resolution, with SoRa 2.8x or 4x magnification. Confocal z-stacks images were maximally projected. For length measurement, cilia ROIs were created by drawing freehand lines along each cilium (ARL13B), and the length of each line was measured using the ROIs. ChP epithelial cilia were manually classified into the following groups, namely (i) Long cilia, (ii) Clustered cilia, and (iii) Scattered cilia, using the previously described categories ^17^.

### CAPture candidate quantification

Confocal Z-stack images were maximally projected and regions containing a cilia-basal body complex cropped from each image (3-5 per image, 5 images per sample). Cropped images of cilia complexes were blinded before analysis. For each cropped image, ROIs were manually drawn for the PCNT-positive area and a similar sized background region. The area, mean PCNT intensity and mean MYH14 intensity was measured for each ROI. MYH14 localisation was calculated as mean intensity of PCNT+ ROI / mean intensity of background ROI. Cilia number was also counted. Values were averaged across all ROIs per sample. For CCDC170, cropped cilia complexes were scored as positive or negative for CCDC170 signal.

### SLC5A5 peak intensity

Confocal z-stacks images were maximally projected. Viewing the DAPI channel, a perpendicular line was drawn across the epithelial layer. The intensity profile along the line was then measured for the SLC5A5 channel and peak value saved. This process was repeated for 10 lines per image and 3-5 images per organoid with the mean peak SLC5A5 calculated per organoid.

### Organoid Cyst Tissue Contrast

Brightfield images acquired on EVOS XL Core microscope were processed on ImageJ/Fiji. ROIs were manually drawn within the organoid cyst area and a similar sized background region. Several ROIs (both within cyst and background) were drawn for each organoid to establish the respective mean brightness intensity. Tissue contrast was calculated as mean brightness intensity within cyst ROIs / mean brightness intensity of background ROIs.

### Nuclei Aspect Ratio

Confocal z-stacks images were maximally projected. DAPI-stained nuclei were segmented and used as reference for two-dimensional nuclear morphometric features. Nuclear aspect ratio (calculated as the ratio of major (a) to minor axis (b) of the nucleus) were measured using the “Analyse particles” function in Fiji. Data points represented were taken from n=3 independent organoid batches from 3 cell lines (H1 and H9-derived for developmental time course analysis; H1 and KOLF2.1J-derived for VPA experiments).

### Statistics

Statistics were performed in R (v 4.4.0) or GraphPad Prism (v10.6.1). For each experiment the n is represented by different batches of organoids.

## Supplementary Figures

**Supplementary figure S1.**
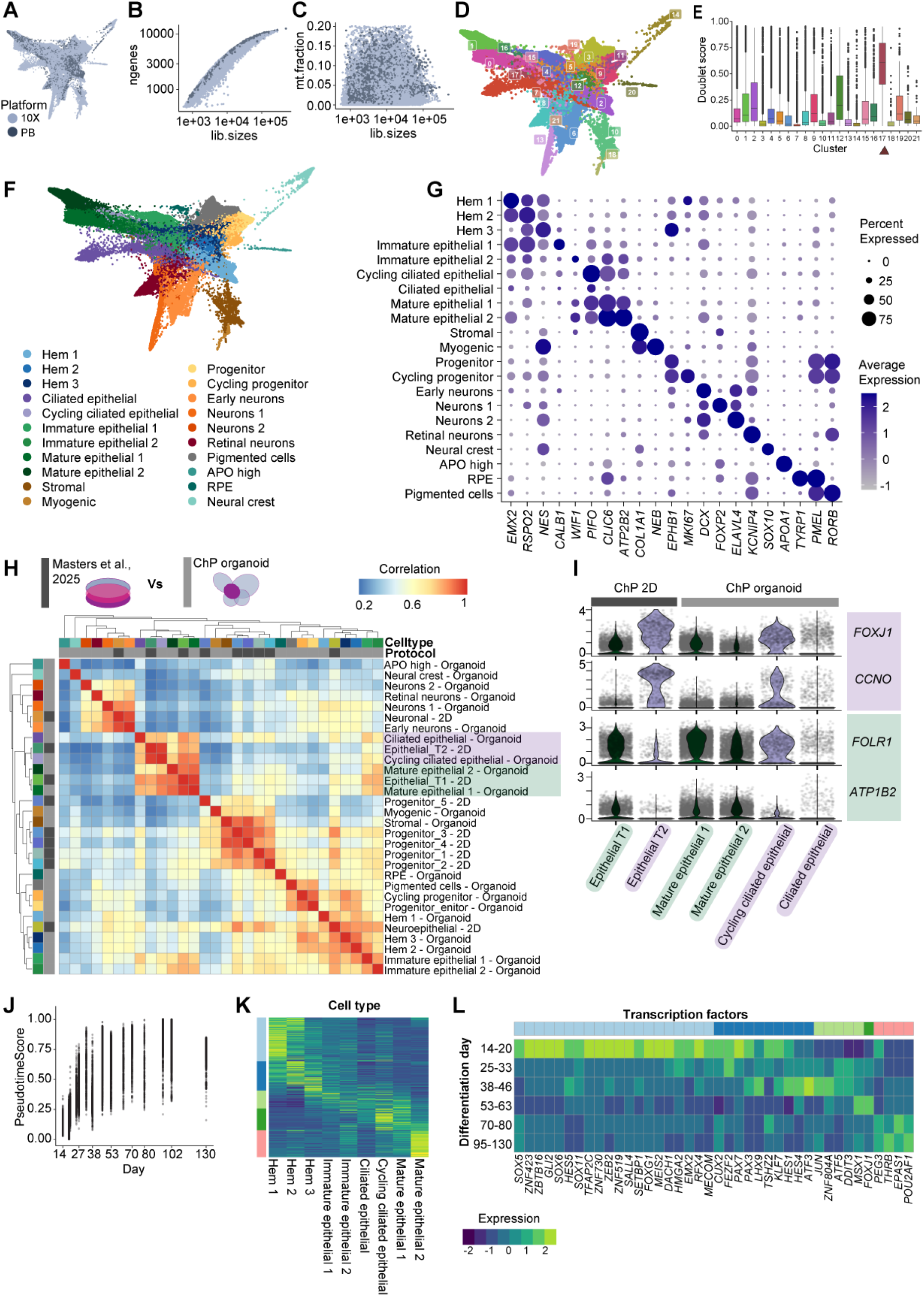
QC of scRNA-seq. **(A)** FA plot coloured by sequencing platform showing integration of sequencing technologies. Points represent cells following QC. **(B)** Number of genes plotted against total reads (library size) per cell, coloured by sequencing platform. Points represent cells following QC. **(C)** Mitochondrial fraction plotted against library size, coloured by sequencing platform. Points represent cells following QC. **(D)** Leiden clusters identified from scVI analysis displayed on FA embedding. **(E)** Doublet score for each leiden cluster. Cells in cluster leiden 17 (arrow) were excluded from downstream analysis due to high doublet content. **(F)** FA plot labelled by fine cell type annotation. **(G)** Dot plot showing expression of key marker genes used for cluster annotation (Table S2). Dot size indicates the percentage of cells expressing gene and colour scale shows the average expression. **(H)** Correlation of broad cell type clusters found in ChP organoids with an alternative 2D ChP differentiation protocol ^17^. Heatmap shows Pearson correlation between pseudobulk gene expression for each cluster, with high similarity in red and low in blue. Clustering by correlation distance results in major groupings: neuronal; epithelial; progenitor (2D) and stromal (organoid); immature epithelial, hem and neuroepithelium. Highlighted populations show separation of cilia-high (2D T2, purple) and secretion-high (2D T1, green) epithelial subtypes. **(I)** Violin plots showing expression of markers distinguishing previously identified ChP epithelial types ^17^. Distinct cilia-high (2D T2; *FOXJ1, CCNO*) and secretory (2D T1; *FOLR1, ATP1B2*) subtypes are present in both differentiation protocols. **(J)** Correlation of pseudotime with real sample ages in epithelial lineage. **(K)** Heatmap showing expression of epithelial lineage modules across cell types. Pseudobulk expression calculated for each cell type cluster. Expression from low (blue) to high (yellow). **(L)** Expression of transcription factors identified in epithelial lineage modules across time in ChP organoids. Pseudobulk expression calculated for all epithelial lineage cells grouped by ages as indicated. Expression from low (blue) to high (yellow).

**Supplementary figure S2.**
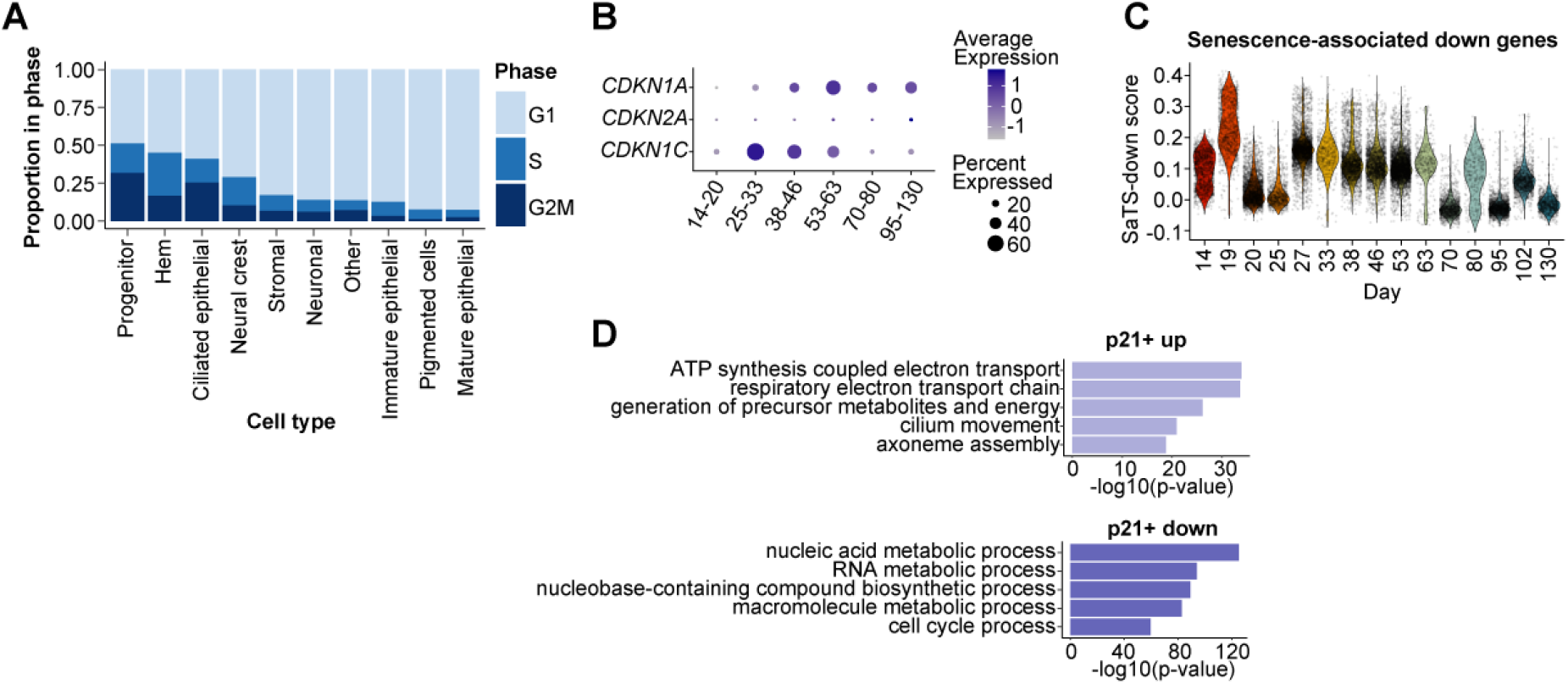
Cell cycle changes across cell types. **(A)** Proportion of cells in each cell cycle phase across broad cell types. **(B)** Expression of selected cyclin-dependent kinase genes across time in the ChP epithelial lineage. Dot size indicates the percentage of cells in each time period expressing the gene and colour scale shows average gene expression. **(C)** Module score for senescence associated transcriptomic signature (SaTS) downregulated genes ^24^ across differentiation time in epithelial lineage cells. **(D)** Pathways differentially expressed in p21+ cells compared to p21-cells within the ChP epithelial lineage.

**Supplementary figure S3.**
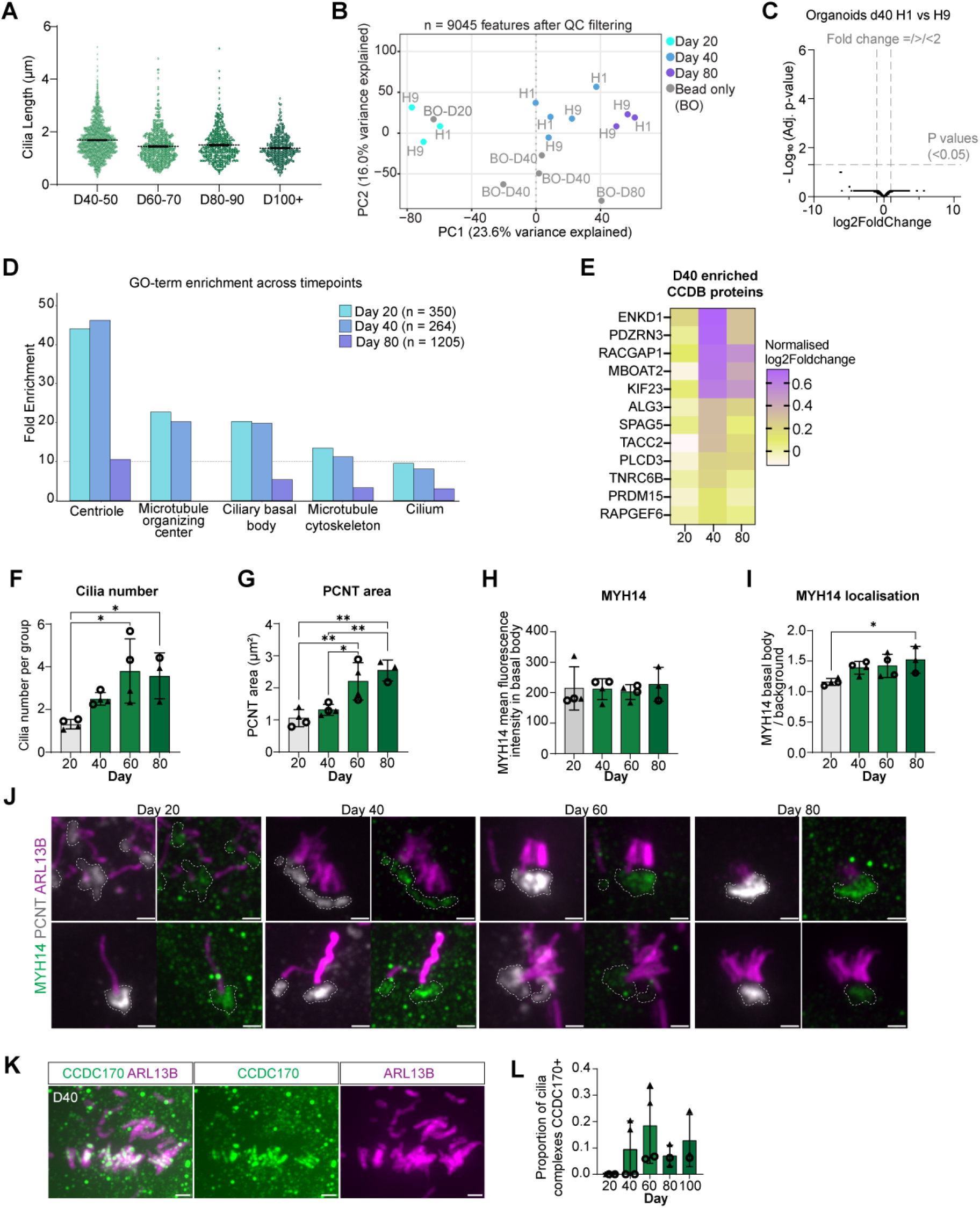
MYH14 as a novel basal body protein in ciliated ChP epithelium. **(A)** Cilia length over time in ChP organoid epithelium. Points represent individual cilia. Cilia measured across organoids from separate differentiations, n = 4-5. **(B)** Principle component plot of CAPture-MS proteomics data showing progressive separation of organoid timepoints along PC1 consistent with differentiation trajectory, as well as PC2, revealing differences between bead only (BO) versus CAPture samples. Point colour indicates sample (timepoint or bead-only), labelled by cell line used. A total of 9,045 protein features passed quality control filtering. **(C)** Volcano plot showing differences between H1 and H9 cell lines at day 40. Axes display log2 fold change and - log10(adjusted P-value). No differences were observed between the H1 and H9 datasets, suggesting that the results are robust across the two independent human embryonic cell line-derived organoids. Differential expression performed using the LIMMA linear model. **(D)** Gene Ontology (GO) term enrichment analysis of proteins identified in isolated centrosome samples. Bar chart shows fold enrichment of the top 7 GO cellular component terms associated with cilia and centrosomes for proteins in isolated centrosome samples at D20, 40 and 80. The GO term of nuclear protein-containing complex (GO:0140513) was used as a negative control. **(E)** Heatmap showing proteins recorded above filtering threshold (log2 fold change compared to bead-only > 2.3) at only D40 and present in CCDB database. Protein level as log2foldchange from bead-only normalised by CEP135, from low (yellow) to high (purple), absent from >= 1 replicate excluded (white). **(F)** Cilia number per cluster across time in ChP organoid. Points represent average cilia number per organoid from different batches, n=3-4. ANOVA and Tukey multiple-comparison test. * p < 0.05. **(G)** Average PCNT-positive basal body area increased across organoid differentiation day. Points represent different differentiation batches, n=3-4. ANOVA and Tukey multiple-comparison test. * p < 0.05, ** p < 0.01. **(H)** Fluorescent intensity of MYH14 in cilia basal bodies (defined by PCNT signal) in ChP epithelium over time. Points represent batches, n=3-4. Difference in mean fluorescent intensity tested by ANOVA, ns. **(I)** Localisation of MYH14 across organoid differentiation day. MYH14 intensity measured in PCNT+ basal body divided by MYH14 intensity in background area. Points calculated from average of ROIs per organoid and represent separate differentiation batches (n = 3-4). ANOVA and Tukey multiple-comparison test. * p < 0.05. **(J)** Additional representative confocal images showing typical range of cilia and basal body complexes across ChP organoid differentiation. Immunostaining with antibodies against PCNT (grey), ARL13B (magenta) and MYH14 (green). PCNT+ basal body regions indicated with dashed lines. Scale bar: 1µm. **(K)** Additional representative image of day 40 organoid immunostained with antibodies against CCDC170 (green) and ARL13B (magenta). Scale bar: 1µm. **(L)** Proportion of cilia complexes that contain CCDC170 signal across differentiation day. Points represent different differentiation batches, n = 3-4. ANOVA test ns. Cell lines are indicated by point shape in plots F-I & L: solid triangle H1 and open circle H9.

**Supplementary figure S4.**
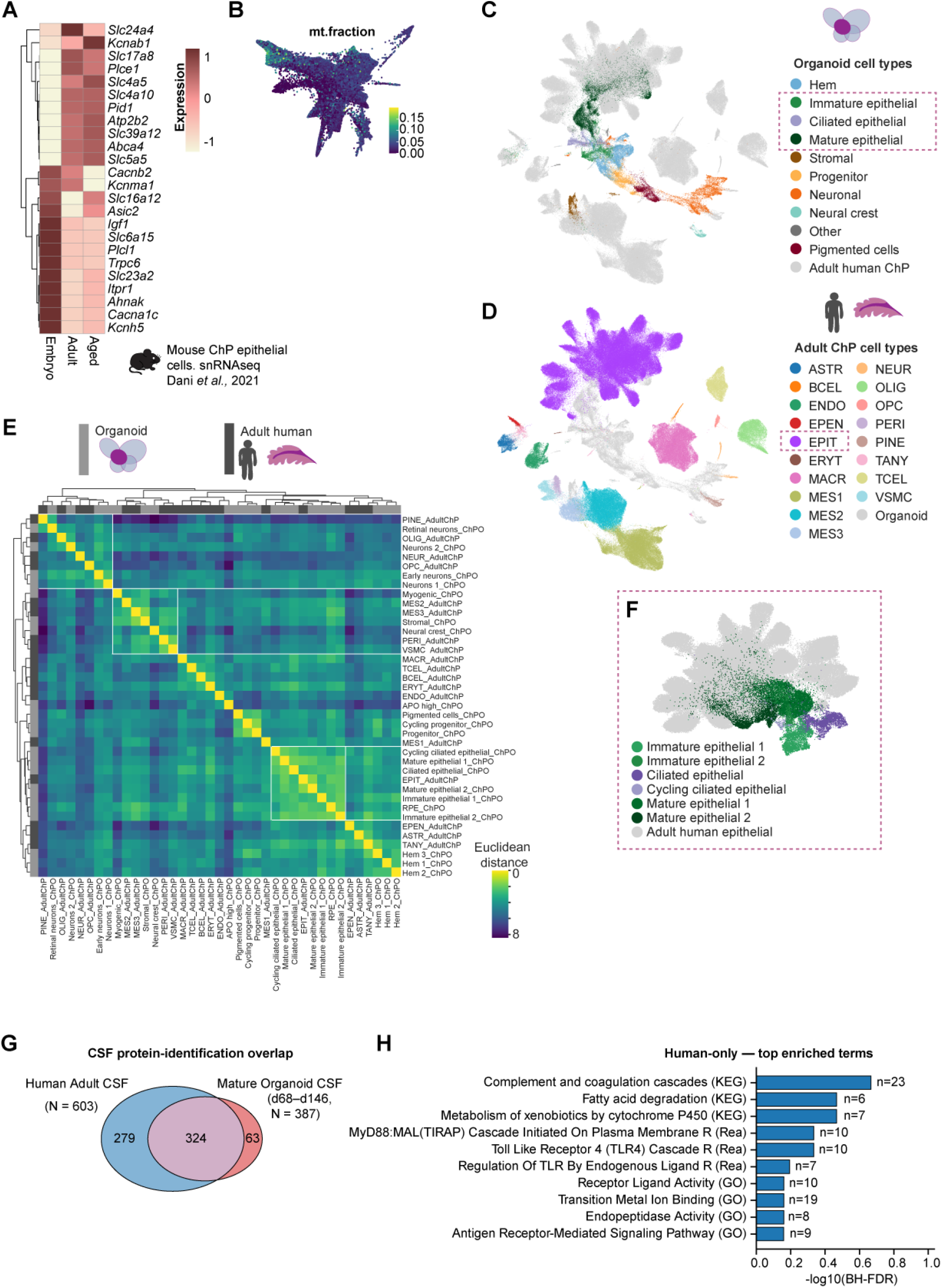
Adult gene expression in organoid, mouse and human ChP. **(A)** Expression of module five transport associated genes across mouse ChP ages (Dani et al., 2021). Pseudobulk expression calculated for all epithelial cells from each age. **(B)** FA plot coloured by mitochondrial fraction highlights high mitochondrial content of mature epithelial cells (top left). Mitochondrial fraction from low (blue) to high (yellow). **(C)** Heatmap showing similarity between organoid and adult human ChP cell types. Adult ChP snRNA sequencing data Mount Sinai cohort. Colour scale represents Euclidean distance between centroids of cell type clusters in scVI latent space, low (yellow) indicates similarity and high (blue) more difference. Integration of datasets shows similarity between organoid and human cell types in groups highlighted by white lines: Neuronal, Mesenchymal / stromal and Epithelial. **(D)** UMAP plot of integrated organoid and adult human ChP datasets coloured by organoid cell types. **(E)** UMAP plot of integrated organoid and adult human ChP datasets coloured by adult cell types. ASTR, astrocyte; BCEL, B cell; ENDO, endothelial; EPEN, ependymal; EPIT, epithelial; ERYT, erythrocyte; MACR, macrophage; MES, mesenchymal; NEUR, neuron; OLIG, oligodendrocyte; PERI, pericyte; PINE, Pinealocyte; TANY, tanycyte; TCEL, T cell; VSMC, vascular smooth muscle cell. **(F)** Venn diagram summarising overlap in identified proteins between ChP organoid ‘CSF’ and adult human CSF. (Table S8). **(G)** Top GO terms for proteins found only in human CSF.

**Supplementary figure S5.**
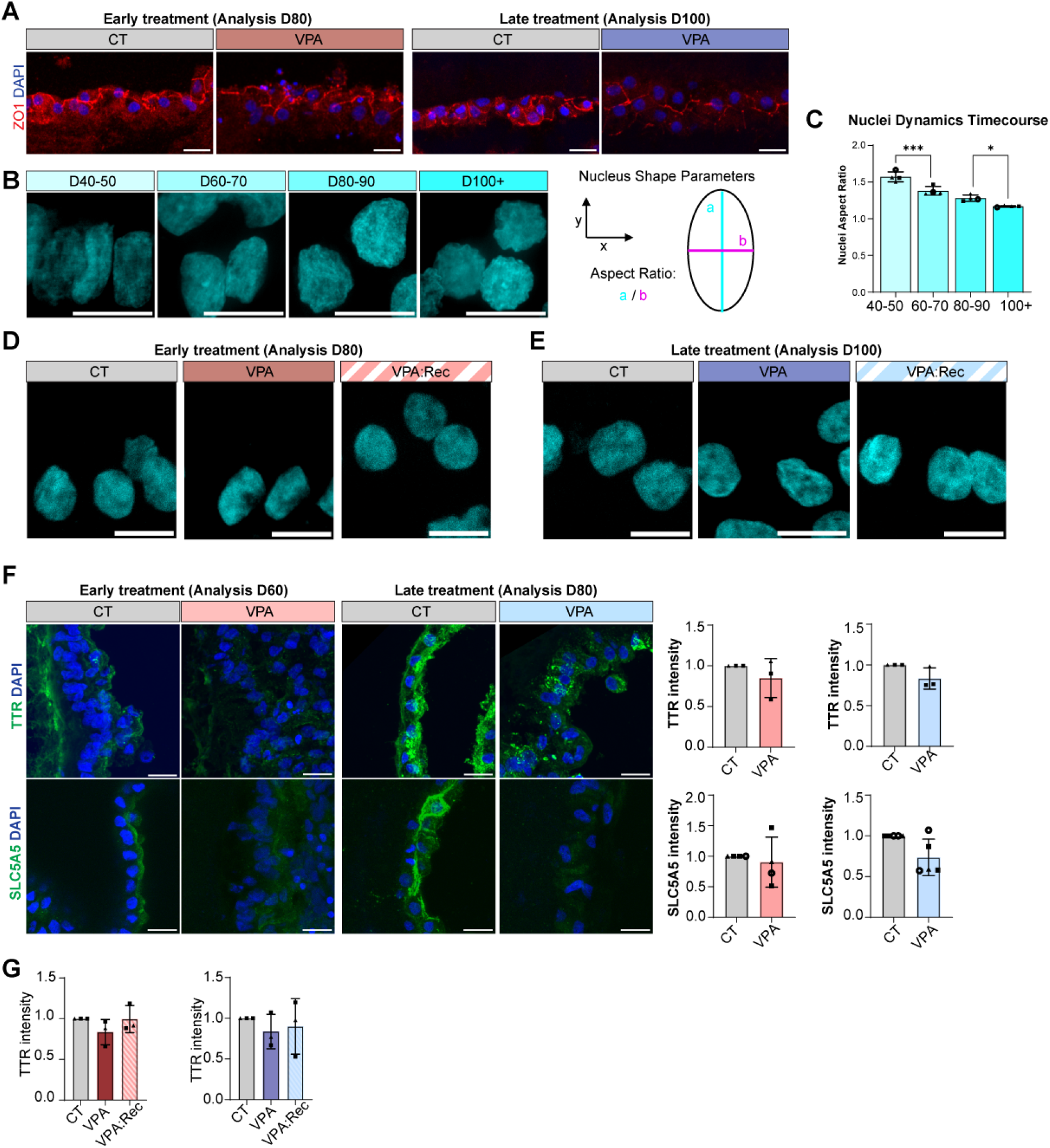
Disruption and recovery following VPA treatment. **(A)** Tight junctions remain intact following VPA-treatment at both timepoints. Immunostaining of D80 (early treatment) and D100 (late treatment) organoids against ZO-1 (red). Nuclei labelled with dapi (blue). Scale bar: 20µm. **(B)** Epithelial nuclei morphology changes over time. Nuclei stained with dapi (cyan). Scale bar: 10µm. **(C)** Nuclear aspect ratio decreases as ChP epithelium over time, consistent with a rounder cell nucleus. Nuclear aspect ratio calculated as major axis (a)/minor axis (b) from maximally projected images. Points represent average from batches, n=4 (with 30-50 nuclei/condition/experiment). One-way ANOVA and Tukey multiple-comparison test * p <0.05, ** p <0.01, *** p <0.001. **(D)** Epithelial nuclei morphology across conditions in early VPA treatments, analysed at D80. Nuclei stained with dapi (cyan). Scale bar: 10µm. **(E)** Epithelial nuclei morphology across conditions in late VPA treatments, analysed at D100. Nuclei stained with dapi (cyan). Scale bar: 10µm. **(F)** VPA treatment for 20 days decreased secretion markers at both treatment timepoints. Organoids immunostained against TTR (top) and SLC5A5 (bottom). Intensity measurements at D60 (early treatment) and D80 (late treatment). Points represent batches, n=3-5. T-test, ns in all cases. **(G)** Intensity of TTR across early (left) and late (right) VPA treatment. Normalised to control. Points indicate batches, n=3. One-way ANOVA, ns. Cell lines are indicated by point shape in plots: solid triangle H1, open circle H9 and solid square KOLF2.1J.

## References

1. Ho, K.H., Candat, A., Scarpetta, V., Brill, M.S., Spassky, N., Patrizi, A., Brill, M.S., and Spassky, N. (2023). Choroid plexuses carry nodal-like cilia that undergo axoneme regression from early adult stage. 1–11. 10.1016/j.devcel.2023.10.003.

2. Lun, M.P., Monuki, E.S., and Lehtinen, M.K. (2015). Development and functions of the choroid plexus-cerebrospinal fluid system. Nat. Rev. Neurosci. 16, 445–457. 10.1038/nrn3921.

3. Saunders, N.R., Dziegielewska, K.M., Fame, R.M., Lehtinen, M.K., and Liddelow, S.A. (2023). the Choroid Plexus: a Missing Link in Our Understanding of Brain Development and Function. Physiol. Rev. 103, 919–956. 10.1152/physrev.00060.2021.

4. Verhaege, D., De Nolf, C., Van Acker, L., Claeys, W., Castelein, J., Van Wonterghem, E., Van Imschoot, G., Dujardin, P., De Spiegelaere, W., Hoste, E., et al. (2026). Base barrier cells provide compartmentalization of choroid plexus, brain and CSF. Nat. Neurosci. 29, 551–566. 10.1038/s41593-025-02188-7.

5. Ghersi-Egea, J.F., Strazielle, N., Catala, M., Silva-Vargas, V., Doetsch, F., and Engelhardt, B. (2018). Molecular anatomy and functions of the choroidal blood-cerebrospinal fluid barrier in health and disease. Acta Neuropathol. 135, 337–361. 10.1007/s00401-018-1807-1.

6. Silva-Vargas, V., Maldonado-Soto, A.R., Mizrak, D., Codega, P., and Doetsch, F. (2016). Age-Dependent Niche Signals from the Choroid Plexus Regulate Adult Neural Stem Cells. Cell Stem Cell 19, 643–652. 10.1016/j.stem.2016.06.013.

7. Tsitsou-Kampeli, A., Suzzi, S., and Schwartz, M. (2024). The immune and metabolic milieu of the choroid plexus as a potential target in brain protection. Preprint at Elsevier Ltd, 10.1016/j.tins.2024.05.010.

8. Xu, H., Lotfy, P., Gelb, S., Pragana, A., Hehnly, C., Byer, L.I.J., Shipley, F.B., Zawadzki, M.E., Cui, J., Deng, L., et al. (2024). The choroid plexus synergizes with immune cells during neuroinflammation. Cell 187, 4946–4963.e17. 10.1016/j.cell.2024.07.002.

9. Bayer, S.A., & A.J. (2007). The Human Brain During the Early First Trimester (1st ed.). CRC Press..

10. Dziegielewska, K.M., Ek, J., Habgood, M.D., and Saunders, N.R. (2001). Development of the choroid plexus. Microsc. Res. Tech. 52, 5–20. 10.1002/1097-0029(20010101)52:1<5::AID-JEMT3>3.0.CO;2-J.

11. Shuangshoti, S., and Netsky, M.G. (1966). Histogenesis of choroid plexus in man. American Journal of Anatomy 118, 283–315. 10.1002/aja.1001180114.

12. Barkho, B.Z., and Monuki, E.S. (2015). Proliferation of cultured mouse choroid plexus epithelial cells. PLoS One 10. 10.1371/journal.pone.0121738.

13. Chouaf-Lakhdar, L., Fèvre-Montange, M., Brisson, C., Strazielle, N., Gamrani, H., and Didier-Bazès, M. (2003). Proliferative activity and nestin expression in periventricular cells of the adult rat brain. Neuroreport 14, 633–636. 10.1097/00001756-200303240-00022.

14. Korzhevskii, D.E. (2000). Proliferative zones in the epithelium of the choroid plexuses of the human embryo brain. Neurosci. Behav. Physiol. 30, 509–512. 10.1007/BF02462607.

15. Liddelow, S.A., Dziegielewska, K.M., VandeBerg, J.L., and Saunders, N.R. (2010). Development of the lateral ventricular choroid plexus in a marsupial, Monodelphis domestica. Cerebrospinal Fluid Res. 7. 10.1186/1743-8454-7-16.

16. Mao, S., Song, R., Jin, S., Pang, S., Jovanovic, A., Zimmerman, A., Li, P., Wu, X., Wendland, M.F., Lin, K., et al. (2025). Multicilia dynamically transduce Sonic Hedgehog signaling to regulate choroid plexus functions. Cell Rep. 44. 10.1016/j.celrep.2025.115383.

17. Masters, H., Wang, S., Tu, C., Nguyen, Q., Sha, Y., Karikomi, M.K., Fung, P.S.R., Tran, B., Martel, C., Kwang, N., et al. (2025). Sequential emergence and contraction of epithelial subtypes in the prenatal human choroid plexus revealed by a stem cell model. Nature Communications 16. 10.1038/s41467-025-60361-9.

18. Pellegrini, L., Bonfio, C., Chadwick, J., Begum, F., Skehel, M., and Lancaster, M.A. (2020). Human CNS barrier-forming organoids with cerebrospinal fluid production. Science (1979). 5626, 1–21.

19. Qiu, F., Huang, Y., Dziegielewska, K.M., Habgood, M.D., and Saunders, N.R. (2025). Gestational Valproate Exposure Induces Tissue-Specific Transcriptomic Changes in the Neonatal Brain and Choroid Plexus in a Rat Model of Epilepsy, GAERS. European Journal of Neuroscience 61. 10.1111/ejn.70139.

20. Dani, N., Herbst, R.H., Habib, N., Head, J., Dionne, D., Nguyen, L., McCabe, C., Cui, J., Shipley, F.B., Jang, A., et al. (2021). A cellular and spatial map of the choroid plexus across brain ventricles and ages. Cell, 1–19. 10.1101/627539.

21. Dohrmann, G.J., and Bucy, P.C. (1970). Human choroid plexus: a light and electron microscopic study. J. Neurosurg. 33, 506–516.

22. Sakaue-Sawano, A., Kurokawa, H., Morimura, T., Hanyu, A., Hama, H., Osawa, H., Kashiwagi, S., Fukami, K., Miyata, T., Miyoshi, H., et al. (2008). Visualizing Spatiotemporal Dynamics of Multicellular Cell-Cycle Progression. Cell 132, 487–498. 10.1016/j.cell.2007.12.033.

23. Dani, N., Herbst, R.H., McCabe, C., Green, G.S., Kaiser, K., Head, J.P., Cui, J., Shipley, F.B., Jang, A., Dionne, D., et al. (2021). A cellular and spatial map of the choroid plexus across brain ventricles and ages. Cell 184, 3056–3074.e21. 10.1016/j.cell.2021.04.003.

24. Watson, L.A., Adelsheim, Z., Carter, M.J., Carter, G.T., Jimenez-Reyes, K.L., Du, H., Zhu, Z., Berry, D.B., Paredez, M.C., Palaniappan, R.H., et al. (2026). Persistent and transient senescent cells contribute to brain-barrier development. Cell. 10.1016/j.cell.2026.05.022.

25. Giovannini, C., Gramantieri, L., Minguzzi, M., Fornari, F., Chieco, P., Grazi, G.L., and Bolondi, L. (2012). CDKN1C/P57 is regulated by the Notch target gene Hes1 and induces senescence in human hepatocellular carcinoma. American Journal of Pathology 181, 413–422. 10.1016/j.ajpath.2012.04.019.

26. Lewis, M., and Stracker, T.H. (2021). Transcriptional regulation of multiciliated cell differentiation. Preprint at Elsevier Ltd, 10.1016/j.semcdb.2020.04.007.

27. Nemajerova, A., Kramer, D., Siller, S.S., Herr, C., Shomroni, O., Pena, T., Suazo, C.G., Glaser, K., Wildung, M., Steffen, H., et al. (2016). TAp73 is a central transcriptional regulator of airway multiciliogenesis. Genes Dev. 30, 1300–1312. 10.1101/gad.279836.116.

28. Carden, S., Vitiello, E., Rosa e Silva, I., Holder, J., Quarantotti, V., Kishore, K., Roamio Franklin, V.N., D’Santos, C., Ochi, T., van Breugel, M., et al. (2023). Proteomic profiling of centrosomes across multiple mammalian cell and tissue types by an affinity capture method. Dev. Cell 58, 2393–2410.e9. 10.1016/j.devcel.2023.09.008.

29. Alves-Cruzeiro, J.M.D.C., Nogales-Cadenas, R., and Pascual-Montano, A.D. (2014). CentrosomeDB: A new generation of the centrosomal proteins database for Human and Drosophila melanogaster. Nucleic Acids Res. 42. 10.1093/nar/gkt1126.

30. Firat-Karalar, E.N., Rauniyar, N., Yates, J.R., and Stearns, T. (2014). Proximity interactions among centrosome components identify regulators of centriole duplication. Current Biology 24, 664–670. 10.1016/j.cub.2014.01.067.

31. Gupta, G.D., Coyaud, É., Gonçalves, J., Mojarad, B.A., Liu, Y., Wu, Q., Gheiratmand, L., Comartin, D., Tkach, J.M., Cheung, S.W.T., et al. (2015). A Dynamic Protein Interaction Landscape of the Human Centrosome-Cilium Interface. Cell 163, 1484–1499. 10.1016/j.cell.2015.10.065.

32. Van Dam, T.J.P., Wheway, G., Slaats, G.G., Huynen, M.A., and Giles, R.H. (2013). The SYSCILIA gold standard (SCGSv1) of known ciliary components and its applications within a systems biology consortium. Preprint, 10.1186/2046-2530-2-7.

33. Nigg, E.A., and Holland, A.J. (2018). Once and only once: Mechanisms of centriole duplication and their deregulation in diseases. Preprint at Nature Publishing Group, 10.1038/nrm.2017.127.

34. Zhang, Y.C., Bai, Y.F., Yuan, J.F., Shen, X.L., Xu, Y.L., Jian, X.X., Li, S., Song, Z.Q., Hu, H. Bin, Li, P.Y., et al. (2021). CEP55 promotes cilia disassembly through stabilizing Aurora a kinase. Journal of Cell Biology 220. 10.1083/JCB.202003149.

35. Rashidieh, B., Shohayeb, B., Bain, A.L., Fortuna, P.R.J., Sinha, D., Burgess, A., Mills, R., Adams, R.C., Lopez, J.A., Blumbergs, P., et al. (2021). Cep55 regulation of PI3K/Akt signaling is required for neocortical development and ciliogenesis. PLoS Genet. 17. 10.1371/journal.pgen.1009334.

36. Yang, Y., Ran, J., Liu, M., Li, D., Li, Y., Shi, X., Meng, D., Pan, J., Ou, G., Aneja, R., et al. (2014). CYLD mediates ciliogenesis in multiple organs by deubiquitinating Cep70 and inactivating HDAC6. Cell Res. 24, 1342–1353. 10.1038/cr.2014.136.

37. Meyer-Miner, A., Van Gennip, J.L.M., Henke, K., Harris, M.P., and Ciruna, B. (2022). Resolving primary pathomechanisms driving idiopathic-like spinal curvature using a new katnb1 scoliosis model. iScience 25. 10.1016/j.isci.2022.105028.

38. Yu, W., Qiang, L., Solowska, J.M., Karabay, A., Korulu, S., and Baas, P.W. (2008). The Microtubule-severing Proteins Spastin and Katanin Participate Differently in the Formation of Axonal Branches. Mol. Biol. Cell 19, 1485–1498. 10.1091/mbc.E07-09.

39. Hiramatsu, K., Nishio, S.Y., Kitajiri, S.I., Kitano, T., Moteki, H., and Usami, S.I. (2021). Prevalence and clinical characteristics of hearing loss caused by myh14 variants. Genes (Basel). 12. 10.3390/genes12101623.

40. Iyadurai, S., Arnold, W.D., Kissel, J.T., Ruhno, C., Mcgovern, V.L., Snyder, P.J., Prior, T.W., Roggenbuck, J., Burghes, A.H., and Kolb, S.J. (2017). Variable phenotypic expression and onset in MYH14 distal hereditary motor neuropathy phenotype in a large, multigenerational North American family. Muscle Nerve 56, 341–345. 10.1002/mus.25491.

41. Donaudy, F., Snoeckx, R., Pfister, M., Zenner, H.-P., Blin, N., Di Stazio, M., Ferrara, A., Lanzara, C., Ficarella, R., Declau, F., et al. (2004). Nonmuscle Myosin Heavy-Chain Gene MYH14 Is Expressed in Cochlea and Mutated in Patients Affected by Autosomal Dominant Hearing Impairment (DFNA4).

42. Hirota, Y., Meunier, A., Huang, S., Shimozawa, T., Yamada, O., Kida, Y.S., Inoue, M., Ito, T., Kato, H., Sakaguchi, M., et al. (2010). Planar polarity of multiciliated ependymal cells involves the anterior migration of basal bodies regulated by non-muscle myosin II. Development 137, 3037–3046. 10.1242/dev.050120.

43. Sigg, M.A., Menchen, T., Lee, C., Johnson, J., Jungnickel, M.K., Choksi, S.P., Garcia, G., Busengdal, H., Dougherty, G.W., Pennekamp, P., et al. (2017). Evolutionary Proteomics Uncovers Ancient Associations of Cilia with Signaling Pathways. Dev. Cell 43, 744–762.e11. 10.1016/j.devcel.2017.11.014.

44. Jiang, P., Li, Y., Poleshko, A., Medvedeva, V., Baulina, N., Zhang, Y., Zhou, Y., Slater, C.M., Pellegrin, T., Wasserman, J., et al. (2017). The Protein Encoded by the CCDC170 Breast Cancer Gene Functions to Organize the Golgi-Microtubule Network. EBioMedicine 22, 28–43. 10.1016/j.ebiom.2017.06.024.

45. Aguilera, P., Hernández-Cruz, A., Franco-Pérez, J., Peralta-Arrieta, I., and Alquisiras-Burgos, I. (2026). The antiaging protein Klotho is a key factor in susceptibility to cerebral ischemia. Preprint at Elsevier Ltd, 10.1016/j.neuroscience.2026.03.005.

46. Kuro-O, M.*, Matsumura, Y., Aizawa, H., Kawaguchi, H., Suga, T., Utsugi, T., Ohyama, Y., Kurabayashi, M., Kaname, T., Kumek, E., et al. (1997). Mutation of the mouse klotho gene leads to a syndrome resembling ageing.

47. Costa-Brito, A.R., Quintela, T., Gonçalves, I., Duarte, A.C., Costa, A.R., Arosa, F.A., Cavaco, J.E., Lemos, M.C., Cecília, &, and Santos, R.A. (2035). The Choroid Plexus Is an Alternative Source of Prolactin to the Rat Brain. 10.1007/s12035-020-02267-9/Published.

48. Savaskan, N.E., Rocha, L., Kotter, M.R., Baer, A., Lubec, G., Van Meeteren, L.A., Kishi, Y., Aoki, J., Moolenaar, W.H., Nitsch, R., et al. (2007). Autotaxin (NPP-2) in the brain: Cell type-specific expression and regulation during development and after neurotrauma. Cellular and Molecular Life Sciences 64, 230–243. 10.1007/s00018-006-6412-0.

49. Patel, B.N., and David, S. (1997). A novel glycosylphosphatidylinositol-anchored form of ceruloplasmin is expressed by mammalian astrocytes. Journal of Biological Chemistry 272, 20185–20190. 10.1074/jbc.272.32.20185.

50. Eikel, D., Lampen, A., and Nau, H. (2006). Teratogenic effects mediated by inhibition of histone deacetylases: Evidence from quantitative structure activity relationships of 20 valproic acid derivatives. Chem. Res. Toxicol. 19, 272–278. 10.1021/tx0502241.

51. Walmod, P.S., Skladchikova, G., Kawa, A., Berezin, V., and Bock, E. (1999). Antiepileptic teratogen valproic acid (VPA) modulates organisation and dynamics of the actin cytoskeleton. Cell Motil. Cytoskeleton 42, 241–255. 10.1002/(SICI)1097-0169(1999)42:3<241::AID-CM7>3.0.CO;2-3.

52. Christensen, J., Grønborg, T.K., Merete, M., Sørensen, J., Schendel, D., Parner, E.T., Pedersen, L.H., and Vestergaard, M. Prenatal Valproate Exposure and Risk of Autism Spectrum Disorders and Childhood Autism.

53. Gottlicher, M., Minucci, S., Zhu, P., Kramer, O.H., Schimpf, A., Giavara, S., Sleeman, J.P., Lo Coco, F., Nervi, C., Pelicci, P.G., et al. (2001). Valproic acid defines a novel class of HDAC inhibitors inducing differentiation of transformed cells.

54. Phiel, C.J., Zhang, F., Huang, E.Y., Guenther, M.G., Lazar, M.A., and Klein, P.S. (2001). Histone Deacetylase is a Direct Target of Valproic Acid, a Potent Anticonvulsant, Mood Stabilizer, and Teratogen. Journal of Biological Chemistry 276, 36734–36741. 10.1074/jbc.M101287200.

55. Wolters, J.E., van Breda, S.G., Claessen, S.M., de Kok, T.M., and Kleinjans, J.C. (2018). Data on novel DNA methylation changes induced by valproic acid in human hepatocytes. Data Brief 16, 161–171. 10.1016/j.dib.2017.11.031.

56. Tanabe, M., Saito, Y., Takasaki, A., Nakano, K., Yamamoto, S., Suzuki, C., Kawamura, N., Hattori, A., Oikawa, M., Nagashima, S., et al. (2025). Role of immature choroid plexus in the pathology of model mice and human iPSC-derived organoids with autism spectrum disorder. Cell Rep. 44. 10.1016/j.celrep.2024.115133.

57. Yang, A.C., Kern, F., Losada, P.M., Agam, M.R., Maat, C.A., Schmartz, G.P., Fehlmann, T., Stein, J.A., Schaum, N., Lee, D.P., et al. (2021). Dysregulation of brain and choroid plexus cell types in severe COVID-19. Nature 595, 565–571. 10.1038/s41586-021-03710-0.

58. Narita, K., and Takeda, S. (2015). Cilia in the choroid plexus: their roles in hydrocephalus and beyond. Front. Cell. Neurosci. 9, 1–5. 10.3389/fncel.2015.00039.

59. Glass, M.R., Matoba, N., Beltran, A.A., Patel, N.K., Farah, T.M., Eswar, K., Bhargava, S., Huang, K., Curtin, I., Ahmed, S., et al. (2026). Human cortical organoids recapitulate inter-individual variability in infant brain-growth trajectories. Cell Stem Cell 33, 142–156.e7. 10.1016/j.stem.2025.12.001.

60. Lennartsson, A., Arner, E., Fagiolini, M., Saxena, A., Andersson, R., Takahashi, H., Noro, Y., Sng, J., Sandelin, A., Hensch, T.K., et al. (2015). Remodeling of retrotransposon elements during epigenetic induction of adult visual cortical plasticity by HDAC inhibitors. Epigenetics Chromatin 8. 10.1186/s13072-015-0043-3.

61. Huangfu, D., Maehr, R., Guo, W., Eijkelenboom, A., Snitow, M., Chen, A.E., and Melton, D.A. (2008). Induction of pluripotent stem cells by defined factors is greatly improved by small-molecule compounds. Nat. Biotechnol. 26, 795–797. 10.1038/nbt1418.

62. Meng, Q., Zhang, W., Wang, X., Jiao, C., Xu, S., Liu, C., Tang, B., and Chen, C. (2022). Human forebrain organoids reveal connections between valproic acid exposure and autism risk. Transl. Psychiatry 12. 10.1038/s41398-022-01898-x.

63. Pellegrini, L., Bonfio, C., Chadwick, J., Begum, F., Skehel, M., and Lancaster, M.A. (2020). Human CNS barrier-forming organoids with cerebrospinal fluid production. Science (1979). 369. 10.1126/science.aaz5626.

## Methods References

1. Lancaster, M.A., and Knoblich, J.A. (2014). Generation of cerebral organoids from human pluripotent stem cells. Nat. Protoc. 9, 2329–2340. 10.1038/nprot.2014.158.

2. Tang See Swee and Apsley, E.J. and P.L. Functional Imaging Methods for Investigating 3D Choroid Plexus Organoids. In (Springer US), pp. 1–18. 10.1007/7651_2024_601.

3. Pellegrini, L., Bonfio, C., Chadwick, J., Begum, F., Skehel, M., and Lancaster, M.A. (2020). Human CNS barrier-forming organoids with cerebrospinal fluid production. Science (1979). 369. 10.1126/science.aaz5626.

4. Lun, A.T.L., Bach, K., and Marioni, J.C. (2016). Pooling across cells to normalize single-cell RNA sequencing data with many zero counts. Genome Biol. 17. 10.1186/s13059-016-0947-7.

5. Germain, P.L., Lun, A., Macnair, W., and Robinson, M.D. (2021). Doublet identification in single-cell sequencing data using scDblFinder. F1000Res. 10, 979. 10.12688/f1000research.73600.1.

6. La Manno, G., Soldatov, R., Zeisel, A., Braun, E., Hochgerner, H., Petukhov, V., Lidschreiber, K., Kastriti, M.E., Lönnerberg, P., Furlan, A., et al. (2018). RNA velocity of single cells. Nature 560, 494–498. 10.1038/s41586-018-0414-6.

7. Traag, V.A., Waltman, L., and van Eck, N.J. (2019). From Louvain to Leiden: guaranteeing well-connected communities. Sci. Rep. 9. 10.1038/s41598-019-41695-z.

8. Dani, N., Herbst, R.H., McCabe, C., Green, G.S., Kaiser, K., Head, J.P., Cui, J., Shipley, F.B., Jang, A., Dionne, D., et al. (2021). A cellular and spatial map of the choroid plexus across brain ventricles and ages. Cell 184, 3056–3074.e21. 10.1016/j.cell.2021.04.003.

9. Masters, H., Wang, S., Tu, C., Nguyen, Q., Sha, Y., Karikomi, M.K., Fung, P.S.R., Tran, B., Martel, C., Kwang, N., et al. (2025). Sequential emergence and contraction of epithelial subtypes in the prenatal human choroid plexus revealed by a stem cell model. Nature Communications 16. 10.1038/s41467-025-60361-9.

10. Dani, N., Herbst, R.H., Habib, N., Head, J., Dionne, D., Nguyen, L., McCabe, C., Cui, J., Shipley, F.B., Jang, A., et al. (2021). A cellular and spatial map of the choroid plexus across brain ventricles and ages. Cell, 1–19. 10.1101/627539.

11. Stoeckius, M., Zheng, S., Houck-Loomis, B., Hao, S., Yeung, B.Z., Mauck, W.M., Smibert, P., and Satija, R. (2018). Cell Hashing with barcoded antibodies enables multiplexing and doublet detection for single cell genomics. Genome Biol. 19. 10.1186/s13059-018-1603-1.

12. Pellegrini, L., Bonfio, C., Chadwick, J., Begum, F., Skehel, M., and Lancaster, M.A. (2020). Human CNS barrier-forming organoids with cerebrospinal fluid production. Science (1979). 5626, 1–21.

13. Carden, S., Vitiello, E., Rosa e Silva, I., Holder, J., Quarantotti, V., Kishore, K., Roamio Franklin, V.N., D’Santos, C., Ochi, T., van Breugel, M., et al. (2023). Proteomic profiling of centrosomes across multiple mammalian cell and tissue types by an affinity capture method. Dev. Cell 58, 2393–2410.e9. 10.1016/j.devcel.2023.09.008.

14. Demichev, V., Messner, C.B., Vernardis, S.I., Lilley, K.S., and Ralser, M. (2020). DIA-NN: neural networks and interference correction enable deep proteome coverage in high throughput. Nat. Methods 17, 41–44. 10.1038/s41592-019-0638-x.

15. Tyanova, S., Temu, T., Sinitcyn, P., Carlson, A., Hein, M.Y., Geiger, T., Mann, M., and Cox, J. (2016). The Perseus computational platform for comprehensive analysis of (prote)omics data. Nat. Methods 13, 731–740. 10.1038/nmeth.3901.

16. Methaneethorn, J. (2018). A systematic review of population pharmacokinetics of valproic acid. Br. J. Clin. Pharmacol. 84, 816–834. 10.1111/bcp.13510.

17. Mao, S., Song, R., Jin, S., Pang, S., Jovanovic, A., Zimmerman, A., Li, P., Wu, X., Wendland, M.F., Lin, K., et al. (2025). Multicilia dynamically transduce Sonic Hedgehog signaling to regulate choroid plexus functions. Cell Rep. 44. 10.1016/j.celrep.2025.115383.

